# *Demuxafy*: Improvement in droplet assignment by integrating multiple single-cell demultiplexing and doublet detection methods

**DOI:** 10.1101/2022.03.07.483367

**Authors:** Drew Neavin, Anne Senabouth, Jimmy Tsz Hang Lee, Aida Ripoll, sc-eQTLGen Consortium, Lude Franke, Shyam Prabhakar, Chun Jimmie Ye, Davis J. McCarthy, Marta Melé, Martin Hemberg, Joseph E. Powell

## Abstract

Recent innovations in droplet-based single-cell RNA-sequencing (scRNA-seq) have provided the technology necessary to investigate biological questions at cellular resolution. With the ability to assay thousands of cells in a single capture, pooling cells from multiple individuals has become a common strategy. Droplets can subsequently be assigned to a specific individual by leveraging their inherent genetic differences, and numerous computational methods have been developed to address this problem. However, another challenge implicit with droplet-based scRNA-seq is the occurrence of doublets - droplets containing two or more cells. The inaccurate assignment of cells to individuals or failure to remove doublets contribute unwanted noise to the data and result in erroneous scientific conclusions. Therefore, it is essential to assign cells to individuals and remove doublets accurately. We present a new framework to improve individual singlet classification and doublet removal through a multi-method intersectional approach.

We developed a framework to evaluate the enhancement in donor assignment and doublet removal through the consensus intersection of multiple demultiplexing and doublet detecting methods. The accuracy was assessed using scRNA-seq data of ∼1.4 million peripheral blood mononucleated cells from 1,034 unrelated individuals and ∼90,000 fibroblast cells from 81 unrelated individuals. We show that our approach significantly improves droplet assignment by separating singlets from doublets and classifying the correct individual compared to any single method. We show that the best combination of techniques varies under different biological and experimental conditions, and we present a framework to optimise cell assignment for a given experiment. We offer *Demuxafy* (https://demultiplexing-doublet-detecting-docs.readthedocs.io/en/latest/index.html) - a framework built-in Singularity to provide clear, consistent documentation of each method and additional tools to simplify and improve demultiplexing and doublet removal. Our results indicate that leveraging multiple demultiplexing and doublet detecting methods improves accuracy and, consequently, downstream analyses in multiplexed scRNA-seq experiments.

## Introduction

Droplet-based single-cell RNA sequencing (scRNA-seq) technologies have provided the necessary tools to profile tens of thousands of single-cell transcriptomes simultaneously^1^. With these technological advances, combining cells from multiple samples in a single capture is now common to increase the sample size while simultaneously reducing batch effects, cost, and time. In addition, following cell capture and sequencing, the droplets can be demultiplexed - each droplet accurately assigned to each individual in the pool^2–5^.

Many scRNA-seq experiments now capture upwards of 20,000 droplets, resulting in ∼16% (3,200) doublets^6^. Current demultiplexing methods can also identify doublets - droplets containing two or more cells - from different individuals (heterogenic doublets). These doublets can significantly alter scientific conclusions if they are not effectively removed. Therefore, it is essential to remove doublets from droplet-based single-cell captures.

However, demultiplexing methods cannot identify droplets containing multiple cells from the same individual (homogenic doublets) and, therefore, cannot identify all doublets in a single capture. If left in the dataset, those doublets could appear as transitional cells between two distinct cell types or a completely new cell type. Accordingly, additional methods have been developed to identify heterotypic doublets (droplets that contain two cells from different cell types) by comparing the transcriptional profile of each droplet to doublets simulated from the dataset^7–13^. It is important to recognise that demultiplexing methods achieve two functions - segregation of cells from different donors and separation of singlets from doublets - while doublet detecting methods solely classify singlets versus doublets.

Therefore, demultiplexing methods and transcription-based doublet detecting methods provide complementary information to improve doublet detection, providing a cleaner dataset and more robust scientific results. There are currently five genetic-based demultiplexing^2–5,14^ and seven transcription-based doublet detecting methods implemented in various languages^7–13^. Under different scenarios, each of these methods is subject to varying performance, and in some instances, biases in their ability to accurately assign cells or detect doublets from certain conditions. The best combination of methods is currently unclear but, undoubtedly, will depend on the dataset and research question.

Therefore, we set out to identify the best combination of genetic-based demultiplexing and transcription-based doublet detecting methods to both correctly remove doublets and partition singlets from different donors. In addition, we have developed a software platform (*Demuxafy*) that performs these intersectional methods and provides additional commands to simplify the execution and interpretation of results for each method (**Figure 1a**).

**Figure 1:**
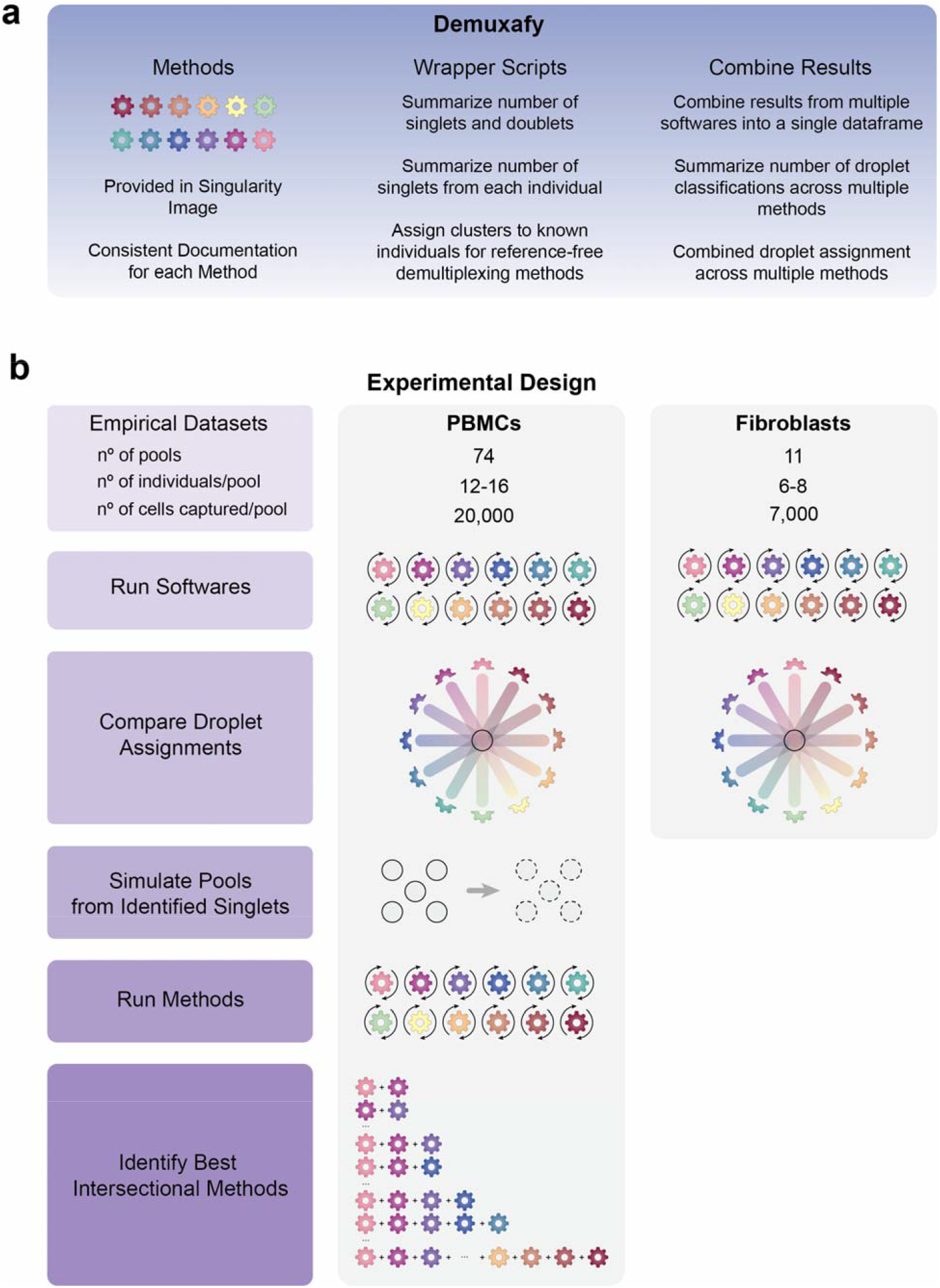
Study design and qualitative method classifications. **a**) Demuxafy is a platform to perform demultiplexing and doublet detecting with consistent documentation. Demuxafy also provides wrapper scripts to quickly summarize the results from each method and assign clusters to each individual with reference genotypes when a reference-free demultiplexing method is used. Finally, Demuxafy provides a script to easily combine the results from multiple different methods into a single data frame and it provides a final assignment for each droplet based on the combination of multiple methods. In addition, Demuxafy provides summaries of the number of droplets classified as singlets or doublets by each method and a summary of the number of droplets assigned to each individual by each of the demultiplexing methods. **b**) Two datasets are included in this analysis - a PBMC dataset and a fibroblast dataset. The PBMC dataset contains 74 pools that captured approximately 20,000 droplets each with 12-16 donor cells multiplexed per pool. The fibroblast dataset contains 11 pools of roughly 7,000 droplets per pool with sizes ranging from six to eight donors per pool. All pools were processed by all demultiplexing and doublet detecting methods and the droplet and donor classifications were compared between the methods and between the PBMCs and fibroblasts. Then the PBMC droplets that were classified as singlets by all methods were taken as ‘true singlets’ and used to generate new pools *in silico*. Those pools were then processed by each of the demultiplexing and doublet detecting methods and intersectional combinations of demultiplexing and doublet detecting methods were tested for different experimental designs.

To compare the demultiplexing and doublet detecting methods, we utilised two large, multiplexed datasets - one that contained ∼1.4 million peripheral blood mononuclear cells (PBMCs) from 1,034 donors and one with ∼94,000 fibroblasts from 81 donors^15^. We used the true singlets from the PBMC dataset to generate new *in silico* pools to assess the performance of each method and the multi-method intersectional combinations (**Figure 1b**).

Here, we compare 12 demultiplexing and doublet detecting methods with different methodological approaches and capabilities and the intersectional combinations. Five of those are demultiplexing methods (*Demuxlet*^*4*^, *Freemuxlet*^*14*^, *ScSplit*^*2*^, *Souporcell*^*5*^, and *Vireo*^*3*^) which leverage the common genetic variation between individuals to identify cells that came from each individual and to identify heterogenic doublets. The seven remaining methods (*DoubletDecon*^*9*^, *DoubletDetection*^*7*^, *DoubletFinder*^*8*^, *ScDblFinder*^*10*^, *Scds*^*11*^, *Scrublet*^*12*^, and *Solo*^*13*^) identify doublets based on their similarity to simulated doublets generated by adding the transcriptional profiles of two randomly selected droplets in the dataset. These methods assume that the proportion of real doublets in the dataset is low, so combining any two droplets is likely to represent the combination of two singlets.

We identify critical differences in the performance of demultiplexing and doublet detecting methods to classify droplets correctly. In the case of the demultiplexing techniques, their performance depends on their ability to identify singlets from doublets and assign a singlet to the correct individual. For doublet detecting methods, the performance is based solely on their ability to differentiate a singlet from a doublet. We identify limitations in identifying specific doublet types and cell types by some methods. In addition, we compare the intersectional combinations of these methods for multiple different experimental designs and demonstrate that intersectional approaches significantly outperform all individual techniques. Thus, the intersectional methods provide enhanced singlet classification and doublet removal - a critical but often under-valued step of droplet-based scRNA-seq processing. Our results demonstrate that intersectional combinations of demultiplexing and doublet detecting software provide significant advantages in droplet-based scRNA-seq preprocessing that can alter results and conclusions drawn from the data. Finally, to provide easy implementation of our intersectional approach, we provide *Demuxafy* (https://demultiplexing-doublet-detecting-docs.readthedocs.io/en/latest/index.html) a complete platform to perform demultiplexing and doublet detecting intersectional methods (**Figure 1a**).

## Results

### Study Design

To study demultiplexing and doublet detecting methods, we developed an experimental design that applies the different techniques to empirical pools and pools generated *in silico* from the combination of true singlets - droplets identified as singlets by every method (**Figure 1a**). For the first phase of this study, we used two empirical multiplexed datasets – the peripheral blood mononuclear cell (PBMC) dataset containing ∼1.4 million cells from 1,034 donors and a fibroblast dataset of ∼94,000 cells from 81 individuals (**Table S1**). We chose these two datasets to assess the methods in heterogeneous (PBMC) and homogeneous (fibroblast) cell types.

### Demultiplexing and Doublet Detecting Methods Perform Similarly for Heterogeneous and Homogeneous Cell Types

We applied the demultiplexing methods (*Demuxlet, Freemuxlet, ScSplit, Souporcell* and *Vireo*) and doublet detecting methods (*DoubletDecon, DoubletDetection, DoubletFinder, ScDblFinder, Scds, Scrublet* and *Solo*) to the two datasets and assessed the results from each method. We first compared the droplet assignments of the different techniques. In the cases where two demultiplexing methods were compared to one another, both the droplet type (singlet or doublet) and the assignment of the droplet to an individual had to match to be considered in agreement. In all other comparisons (*i*.*e*., demultiplexing versus doublet detecting and doublet detecting versus doublet), only the droplet type (singlet or doublet) was considered for agreement. We found that the two method types were more similar to other methods of the same type (*i*.*e*., demultiplexing versus demultiplexing and doublet detecting versus doublet detecting) than they were to methods from a different type (demultiplexing methods versus doublet detecting methods (**Figure 2a-b**). We found that the similarity of the demultiplexing and doublet detecting methods to one another was consistent in the PBMC and fibroblast datasets (Pearson correlation R = 0.78, *P*-value = 8.1*10^−28^; **Figure 2a-b, Figure S1a**). In addition, demultiplexing methods were more similar than doublet detecting methods for both the PBMC and fibroblast datasets (Wilcoxon rank-sum test: *P* < 0.01; **Figure 2a-b** and **S1**).

**Figure 2:**
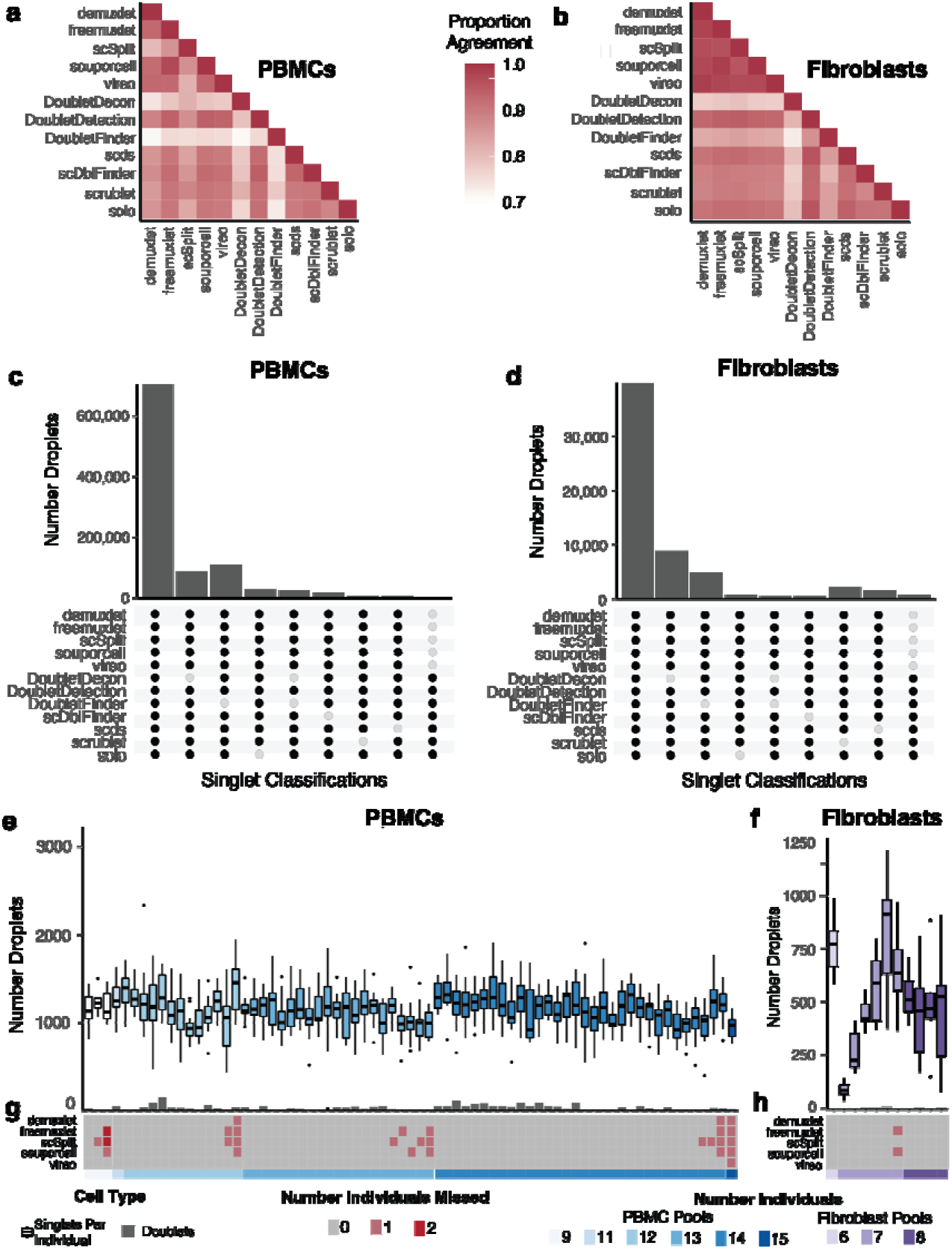
Demultiplexing and Doublet Detecting Method Performance Comparison. **a-b)** Heatmap of agreement of droplet classifications between different methods for the PBMCs (**a**) and fibroblasts (**b**). **c-d)** Upset plot of the PBMC (**c**) and fibroblast (**d**) droplets classified as singlets by different methods. The majority of droplets are classified as singlets by all methods, but there are small numbers of droplets classified as doublets by specific methods. **e-f**) The number of droplets classified as singlets (box plots) and doublets (bar plots) by all methods in the PBMC (**e**) and fibroblast (**f**) pools. **g-h**) The number of donors that were not identified by each method in each pool for PBMCs (**g**) and fibroblasts (**h**). PBMC: peripheral blood mononuclear cell.

The number of droplets classified as singlets by multiple methods and the QC metrics for each grouping of methods was consistent for both the PBMC and fibroblast datasets (**Figures 2c-d, S2**). These data indicate that the methods behave similarly, relative to one another, for heterogeneous and homogeneous datasets.

Next, we sought to identify the droplets concordantly classified by all demultiplexing and doublet detecting methods in the PBMC and fibroblast datasets. On average, 1,146 singlets were identified for each individual by all the methods in the PBMC dataset. Likewise, 504 droplets were identified as singlets for each individual by all the methods in the fibroblast pools. However, the concordance of doublets identified by all methods was very low for both datasets (**Figure 2e-f**). Notably, while the concordance between the two approaches could be high (**Figure 2a-b**), the consistency of classifying a droplet as a doublet by all methods was relatively low (**Figure 2e-f**). This suggests that doublet identification is not consistent between all the methods. Therefore, further investigation is required to identify the reasons for these inconsistencies between methods. It also suggests that combining multiple methods for doublet classification may be necessary for more complete doublet removal. Further, some methods could not identify all the individuals in each pool (**Figure 2g-h**). The non-concordance between different methods demonstrates the need to effectively test each method on a dataset where the droplet types are known.

### Computational Resources Vary for Demultiplexing and Doublet Detecting Methods

We recorded each method’s computational resources for the PBMC pools, with ∼20,000 cells captured per pool (**Table S1**). *ScSplit* took the most time and steps to run the demultiplexing methods, but *Demuxlet* and *Freemuxlet* used the most memory. Solo took the longest time, and most memory to run for the Doublet Detecting methods but is the only method built to be run directly from the command line, making it easy to implement (**Figure S3**).

### Generate Pools with Known Singlets and Doublets

However, there is no gold standard to identify which droplets are singlets or doublets. Therefore, in the second phase of our experimental design (**Figure 1a**), we used the PBMC droplets classified as singlets by all methods to generate new pools *in silico*. We chose to use the PBMC dataset since our first analyses indicated that method performance is similar for homogeneous (fibroblast) and heterogeneous (PBMC) cell types (**Figure 2** and **S1**) and because we had many more individuals available to generate new pools from the PBMC dataset (**Table S1**).

We generated 70 pools - ten each of pools that included two, four, eight, 16, 32, 64 or 128 individuals (**Table S2**). We assume a maximum 20% doublet rate as it is unlikely researchers would use a technology that has a higher doublet rate (**Figure 3a**).

**Figure 3:**
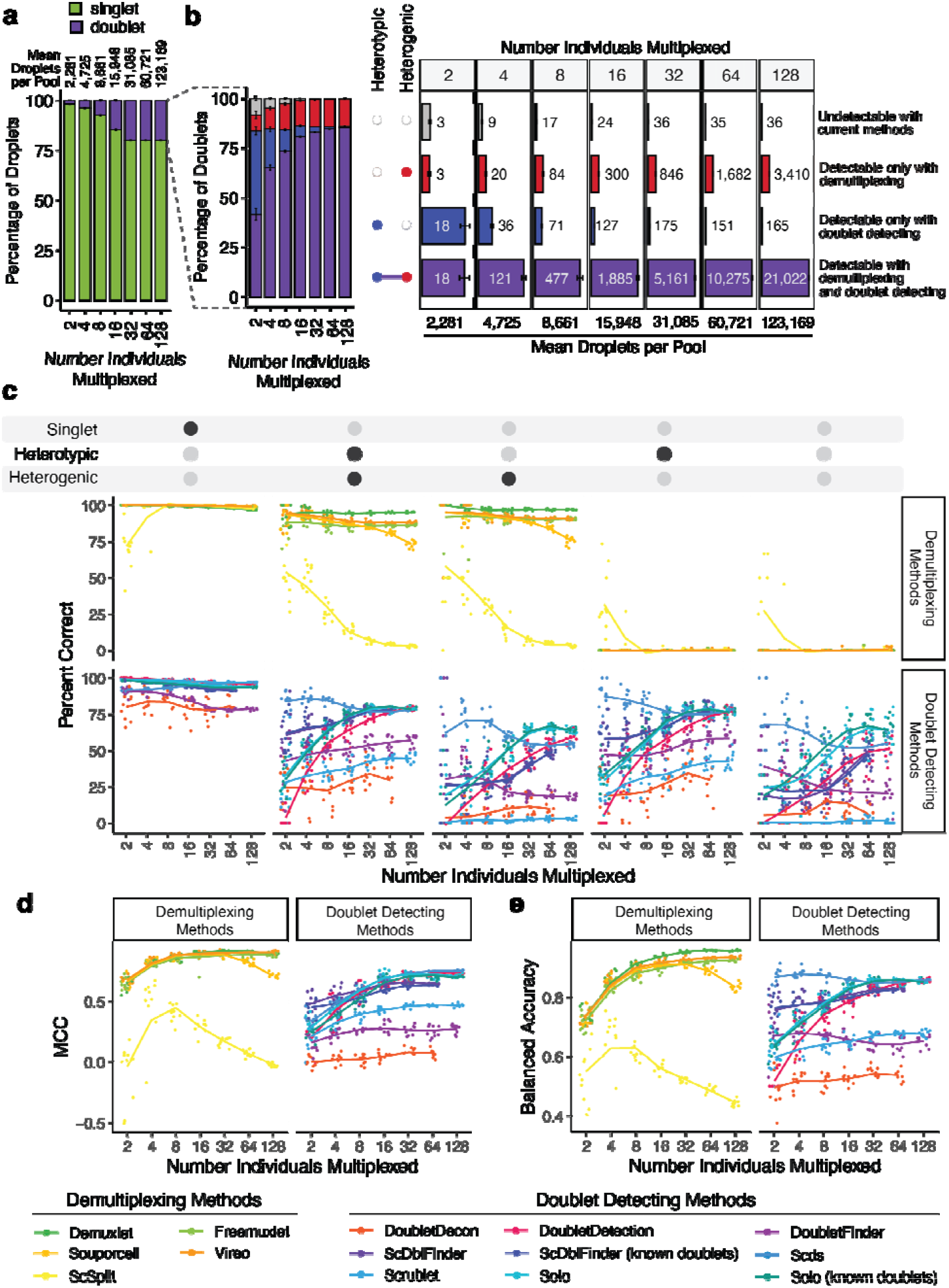
*In silico* Pool Doublet Annotation and Method Performance. **a)** The percent of singlets and doublets in the *in-silico* pools - separated by the number of multiplexed individuals per pool. **b)** The percentage and number of doublets that are heterogenic (detectable by demultiplexing methods), heterotypic (detectable by doublet detecting methods), both (detectable by either method category) and neither (not detectable with current methods) for each multiplexed pool size. **c**) Percent of droplets that each of the demultiplexing and doublet detecting methods classified correctly for singlets and doublet subtypes for different multiplexed pool sizes. **d**) Mathew’s Correlation Coefficient (MCC) for each of the methods for each of the multiplexed pool sizes. **e**) Balanced accuracy for each of the methods for each of the multiplexed pool sizes

We next classified the PBMC cell types for each droplet used to generate the *in-silico* pools with Azimuth to quickly identify the heterotypic doublets in the *in-silico* data^16^ (**Figure S4**). As these pools have been generated *in silico* using empirical singlets that have been well annotated, we next identified the proportion of doublets in each pool that were heterogenic, heterotypic, both and neither. This approach demonstrates that a significant percentage of doublets are only detectable by doublet detecting methods (homogenic and heterotypic) for pools with 16 or fewer donors multiplexed (**Figure 3b**).

While the total number of doublets that would be missed if only using demultiplexing methods appears small for fewer multiplexed individuals (**Figure 3b**), it is important to recognise that this is partly a function of the ∼1,146 singlet cells per individual used to generate these pools. Hence, the *in-silico* pools with fewer individuals also have fewer cells. Therefore, to obtain numbers of doublets that are directly comparable to one another, we calculated the number of each doublet type that would be expected to be captured with 20,000 cells when two, four, eight, 16 or 32 individuals were multiplexed (**Figure S5**). These results demonstrate that many doublets would be falsely classified as singlets since they are homogenic when just using demultiplexing methods for a pool of 20,000 cells captured with a 16% doublet rate (**Figure S5**). However, as more individuals are multiplexed, the number of droplets that would not be detectable by demultiplexing methods (homogenic) decreases. This suggests that typical workflows that use only one demultiplexing method to remove doublets from pools that capture 20,000 droplets with 16 or fewer multiplexed individuals fail to adequately remove between 173 (16 multiplexed individuals) and 1,325 (2 multiplexed individuals) doublets that are homogenic and heterotypic which could be detected by doublet detecting methods (**Figure S5**). Therefore, a technique that uses both demultiplexing and doublet detecting methods in parallel will complement more complete doublet removal methods. Consequently, we next set up to identify the demultiplexing and doublet detecting methods that perform the best on their own and in concert with other methods.

### Doublet and Singlet Droplet Classification Effectiveness Varies for Demultiplexing and Doublet Detecting Methods

#### Demultiplexing Methods Fail to Classify Homogenic Doublets

We next investigated what percentage of the droplets were correctly classified by each demultiplexing and doublet detecting method. Demultiplexing methods correctly classify a large portion of the singlets and heterogenic doublets (**Figure 3c**). This pattern is highly consistent across different cell types, with the notable exceptions being decreased correct classifications for erythrocytes and platelets when greater than 16 individuals are multiplexed (**Figure S6)**.

However, *Demuxlet* consistently demonstrates the highest correct heterogenic doublet classification. Further, the percentage of the heterogenic doublets classified correctly by *Souporcell* decreases when large numbers of donors are multiplexed. *ScSplit* is not as effective as the other demultiplexing methods at classifying heterogenic doublets, partly due to the unique doublet classification method, which assumes that the doublets will generate a single cluster separate from the donors (**Table 1**). In addition, we note that all the demultiplexing methods except *ScSplit* are significantly better at detecting heterogenic doublets that are also heterotypic compared to those that are homotypic (**Figure S7**). This may be because reads from two different cells in a single droplet that overlap a given genetic variant are more likely when the two cells are the same cell type. However, importantly, the demultiplexing methods identify almost none of the homogenic doublets for any multiplexed pool size - demonstrating the need to include doublet detecting methods to supplement the demultiplexing method doublet detection.

#### Doublet Detecting Method Classification Performances Vary Greatly

In addition to assessing each of the methods with default settings, we also evaluated *ScDblFinder* and *Solo* with ‘known doublets’ provided. These two methods can take already known doublets and use them when detecting doublets. For these cases, we used the droplets that were classified as doublets by all the demultiplexing methods as ‘known doublets’.

Generally, the doublet detecting methods showed more variation in the percentage of droplets that they classified correctly (*F*-test *P* < 0.04) except for pools that included two or four multiplexed individuals (*F*-test *P* > 0.12). Most of the methods classified a similarly high percentage of singlets correctly, with the exceptions of *DoubletDecon* and *DoubletFinder* for all pool sizes as well as *Scds* for pools containing less than eight individuals (**Figure 3c**). However, unlike the demultiplexing methods, there are explicit cell-type-specific biases for many of the doublet detecting methods (**Figure S8**). These differences are most notable for cell types with fewer cells (*i*.*e*., ASDC and cDC2) and proliferating cells (*i*.*e*., CD4 Proliferating, CD8 Proliferating and NK Proliferating). DoubletDetection and Scrublet preserve the highest percentage of singlets for all proliferating cell types, which may be crucial for specific experimental questions (**Figure S8**).

As expected, all doublet detecting methods identified heterotypic doublets more effectively than homotypic doublets (**Figure 3c**). However, *Scds* classified the most doublets correctly across all doublet types for 16 individuals or fewer pools. Solo was as good or more effective at identifying doublets than *Scds* for pools containing more than 16 individuals. *ScDblFinder* is also among the methods that correctly identifies the highest percentage of doublets, although it performs better for heterotypic doublets than homotypic doublets. It is also important to note that it was not feasible to run *ScDblFinder* or *DoubletDecon* for the largest pools containing 128 multiplexed individuals and an average of 123,169 droplets (range: 119,942 - 127,173 droplets). *ScDblFinder* and *Solo* performed similarly when executed with and without known doublets (79% of *Solo* and 78% of *ScDblFinder P* > 0.05). Further, for the few conditions where the performance of *ScDblFinder* and *Solo* were significantly different with and without using known doublets, the method run without known doublets identified a substantially higher percentage correct than the method run with known doublets (100% of *ScDblFinder* and 96% of *Solo*). This suggests that providing known doublets to *Solo* and *ScDblFinder* does not offer an added benefit.

#### False Singlets and Doublets Demonstrate Different Metrics than Correctly Classified Droplets

We next asked whether specific cell metrics might contribute to false singlet and doublet classifications for different methods. Therefore, we compared the number of genes, number of UMIs, mitochondrial percentage and ribosomal percentage of the false singlets and doublets to equal numbers of correctly classified cells for each demultiplexing and doublet detecting method.

The number of UMIs (**Figure S9** and **Table S3**) and genes (**Figure S10** and **Table S4**) demonstrated very similar distributions for all comparisons and all methods (Spearman ρ = 0.99, *P* < 2.2*10^−308^). The number of UMIs and genes were consistently higher in false singlets and lower in false doublets for most demultiplexing methods except smaller pool sizes and most *Vireo* pools (**Figures S9a** and **S10a**; **Tables S3** and **S4**). The number of UMIs and genes was consistently higher in droplets falsely classified as singlets by the doublet detecting methods than the correctly identified droplets (**Figure S9b** and **S10b**; **Tables S3** and **S4**). However, there was less consistency in the number of UMIs and genes detected in false singlets than correctly classified droplets between the different doublet detecting methods (**Figures S9b** and **S10b**; **Tables S3** and **S4**).

The ribosomal percentage of the droplets falsely classified as singlets or doublets is similar to the correctly classified droplets for most methods - although they are statistically different for larger pool sizes (**Figure S11a** and **Table S5**). However, the false doublets classified by some doublet detecting methods (*DoubletDetection, DoubletFinder, ScDblFinder, ScDblFinder*) with known doublets and *Scds)* demonstrated lower ribosomal percentages (**Figure S11b** and **Table S5**).

Like the ribosomal percentage, the mitochondrial percentage is also relatively similar for false singlets compared to correctly classified droplets for both demultiplexing (**Figure S12a** and **Table S6**) and doublet detecting methods (**Figure S12b**). Still, it is statistically different for larger pool sizes of some techniques (**Figure S12b** and **Table S6**). However, the mitochondrial percentage for false doublets is statistically higher than the correctly classified droplets for most demultiplexing and doublet detecting methods. Still, it is especially noticeable for *Souporcell, Vireo, DoubletDecon* and *DoubletFinder* (**Figure S12b**).

Overall, these results demonstrate a strong relationship between the number of genes and UMIs and limited influence of ribosomal or mitochondrial percentage in a droplet and false classification, suggesting that the number of genes and UMIs can significantly bias singlet and doublet classification by demultiplexing and doublet detecting methods.

#### Performances Vary Between Demultiplexing and Doublet Detecting Method and Across the Number of Multiplexed Individuals

We assessed the method performance with two metrics: the balanced accuracy and the Mathews correlation coefficient (MCC). We chose the balanced accuracy since, with unbalanced group sizes, it is a better measure of performance than accuracy itself. Further, the MCC has been demonstrated as a more reliable statistical measure of performance since it considers all possible categories - true singlets (true positives), false singlets (false positives), true doublets (true negatives) and false doublets (false negatives). Therefore, a high score on the MCC scale indicates high performance in each metric. However, we provide a wide range of performance metrics for each method (**Table S7**). For demultiplexing methods, both the droplet type (singlet or doublet) and the individual assignment were required to be considered a ‘true singlet’. In contrast, only the droplet type (singlet or doublet) was needed for doublet detection methods.

The MCC and balanced accuracy metrics are strikingly similar (Spearman’s ρ = 0.93; *P* < 2.2*10^−16^). The demultiplexing methods (except *ScSplit*) perform better on average than the doublet detecting methods for both the MCC (Student’s *t*-test *P* < 4.4*10^−14_^ and balanced accuracy) and balanced accuracy (Student’s *t*-test *P* < 4*10^−6^). Further, the performance of *Souporcell* decreases for pools with more than 32 individuals multiplexed for both metrics (Student’s *t*-test for MCC: *P* < 7*10^−6^ and balanced accuracy: *P* < 3.2*10^−5^). *Scds, Solo, ScDblFinder* and *DoubletDetection* are among the top-performing doublet detecting methods. Still, a large variation in the MCC and balanced accuracy is observed in smaller pool sizes (Spearman’s *P* = 2.8*10^−3^ for MCC and *P* = 4*10^−4^ for balanced accuracy; **Figure 3d-e**).

Overall, between one and 59% of droplets were incorrectly classified by the demultiplexing or doublet detecting methods depending on the technique and the multiplexed pool size (**Figure S13**). *Demuxlet, Freemuxlet, Souporcell* and *Vireo* demonstrated the lowest percentage of incorrect droplets with about one per cent wrong in the smaller pools (2 multiplexed individuals) and about three per cent inaccurate for pools with at least 16 multiplexed individuals (although *Souporcell* identified a slightly higher per cent of droplets incorrectly in the largest pools). Seeing as some transitional states and cell types are present in low percentages in total cell populations (*i*.*e*., ASDCs at 0.02%), incorrect classification of droplets could alter scientific interpretations of the data, and it is, therefore, ideal for decreasing the number of erroneous assignments as much as possible.

Our results demonstrate significant differences in overall performance between different demultiplexing and doublet detecting methods. We further noticed some differences in the use of the methods. Therefore, we have accumulated these results and each method’s unique characteristics and benefits in a heatmap for visual interpretation (**Figure 4**).

**Figure 4:**
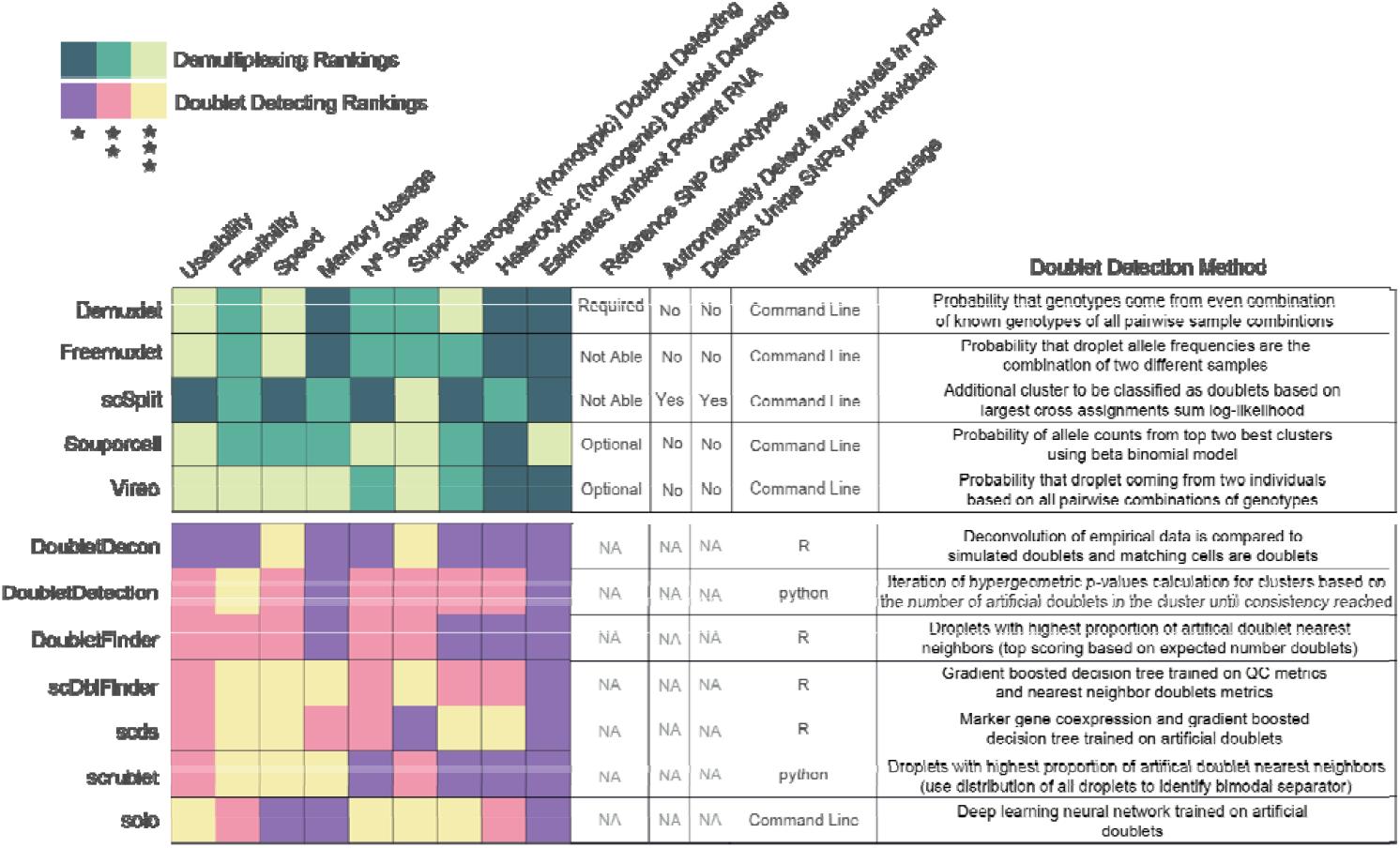
Assessment of each of the demultiplexing and doublet detecting methods. Assessments of a variety of metrics for each of the demultiplexing (top) and doublet detecting (bottom) methods.

### Framework for Improving Singlet Classifications via Method Combinations

After identifying the demultiplexing and doublet detecting methods that performed well individually, we next sought to test whether using intersectional combinations of multiple methods would enhance droplet classifications and provide a software platform - *Demuxafy* - capable of supporting the execution of these intersectional combinations.

We recognise that different experimental designs will be required for other projects. As such, we considered this when testing combinations of methods. We regarded as multiple experiment designs and provided recommendations on two different levels of filtering doublets: 1) a balanced approach that attempts to remove true doublets while not removing too many true singlets (assessed with the MCC) and 2) an approach that removes as many doublets as possible even if some droplets that are true singlets are classified as doublets (assessed with the positive predictive value [PPV]). We considered all possible combinations of methods that achieved greater than 0.5 MCC or greater than 0.8 balanced accuracies for any pool size (**Figure 3d-e**). Those methods included four demultiplexing methods (*Demuxlet, Freemuxlet, Vireo* and *Souporcell*) and four doublet detecting methods (*DoubletDetection, ScDblFinder, Scds* and *Solo*). We also considered two different intersectional methods: 1) more than half had to classify a droplet as a singlet to be called a singlet, and 2) at least half of the methods had to classify a droplet as a singlet to be called a singlet. Significantly, these two intersectional methods only differ when an even number of methods are being considered. For combinations that include demultiplexing methods, the individual called by the majority of the methods is the individual used for that droplet. When ties occur, the individual is considered ‘unassigned’.

#### Combining Multiple Doublet Detecting Methods Improve Doublet Removal for Non-Multiplexed Experimental Designs

For the non-multiplexed experimental design, we considered all possible method combinations of *DoubletDetection, ScDblFinder, Scds* and *Solo* (**Table S9**). We identified important differences depending on the number of droplets captured and have provided recommendations accordingly. We identified that ScDblFinder and Scds is the ideal combination for balanced droplet calling when less than 3,000 droplets are captured. ScDblFinder, *Scds, Solo* and *DoubletDetection* is the best combination when 3,000-10,000 droplets are captured. Scds, *Solo*, and *DoubletDetection* is the best combination when more than 10,000 droplets are captured. It’s important to note that even a slight increase in the MCC significantly impacts the number of true singlets and true doublets classified with the degree of benefit highly dependent on the original method performance (**Figure S14**). The combined method increases the MCC compared to individual doublet detecting methods on average by 0.23 and up to 0.73 - a significant improvement in the MCC (*t*-test FDR < 0.05 for 95% of comparisons). For all combinations, the intersectional droplet method requires more than half of the methods to consider the droplet a singlet to classify it as a singlet (**Figure 5**).

**Figure 5:**
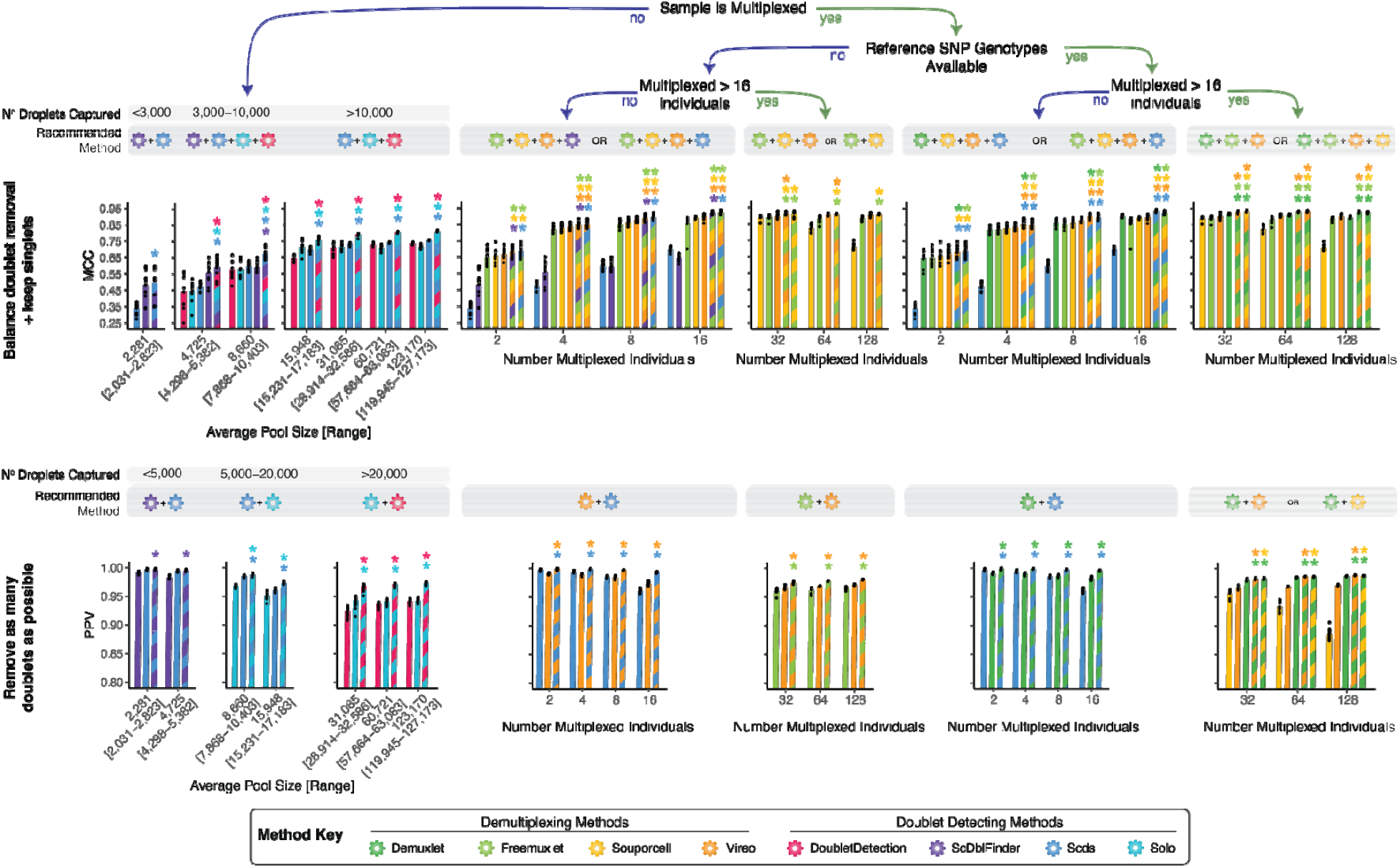
Recommended Method Combinations Dependent on Experimental Design. Method combinations are provided for two different doublet removal levels - a balanced approach that removes doublets and tries to limit the number of singlets that are removed (top panel) and a strict doublet removal approach that eliminates as many doublets as possible even if some singlets are removed (bottom panel). Recommendations are provided for different experimental designs, including those that are not multiplexed (left) and multiplexed (right), including experiments that have reference SNP genotypes available vs those that do not and finally, multiplexed experiments with different numbers of individuals multiplexed. The single-colour bar provides the metrics of individual methods. In contrast, the multi-colour bars provide the metrics for combining techniques as indicated by the colours in those bars. A student’s *t*-test was used to compare the single methods to the combined methods, and significant differences (*P* < 0.05) are indicated with an Asterix (*) and the colour shows the single method that was compared to that combination of methods

For experimental questions where it is crucial to remove as many doublets as possible, the combination of *ScDblFinder* and *Scds* is the best for pools with less than 5,000 droplets captured, *Solo* and *Scds* are ideal when 5,000-10,000 droplets are captured, and the combination of *Solo* and *DoubletDetection* is ideal when more than 10,000 droplets are captured. Notably, the combined method demonstrates a higher PPV than any individual method - a PPV increase of up to 0.16 and an average increase of 0.04 - which is significant for most method comparisons (96.7% of comparisons *t*-test FDR < 0.05). Of note, even a relatively small change in the PPV of 0.02 for a pool that captures 8,661 droplets results in the reannotation of 868 droplets (10% of total droplets). Compared to the individual methods, these intersectional methods decrease the number of true singlets but increase the number of true doublets annotated (**Figure S15**). Again, in all cases, the best intersectional approach is to call a singlet where more than half of the methods classify the droplet as a singlet (**Figure 5**).

#### Combining Multiple Demultiplexing and Doublet Detecting Methods Improve Doublet Removal for Multiplexed Experimental Designs

For experiments where 16 or fewer individuals are multiplexed with reference SNP genotypes available, we considered all possible combinations between *Demuxlet, Freemuxlet, Souporcell, Vireo, DoubletDetection, scDblFinder, Scds* and *Solo* (**Table S10**). To provide a balance between doublet removal and maintaining true singlets, the best combinations are *Demuxlet, Souporcell, Vireo* and *Scds* or *Freemuxlet, Souporcell, Vireo* and *Scds* (**Figure 5**). These intersectional methods increase the MCC compared to the individual methods (*t*-test FDR < 0.05 for 96.4% of comparisons), generally resulting in increased true singlets and doublets compared to the individual methods (**Figure S16**). The improvement in MCC depends on every single method’s performance but, on average, increases by 0.33 (0.14 for demultiplexing methods and 0.43 for doublet detecting methods) and up to 0.89. For experiments where the reference SNP genotypes are unknown, the individuals multiplexed in the pool with 16 or fewer individuals multiplexed, *Freemuxlet, Souporcell, Vireo* and *ScDblFinder* or *Freemuxlet, Souporcell, Vireo* and *Scds* are the ideal methods (**Figure 5**). These intersectional methods again significantly increase the MCC on average by 0.35 (0.16 for demultiplexing methods and 0.43 for doublet detecting methods) compared to any of the individual techniques that could be used for this experimental design (*t*-test FDR < 0.05 for 94.2% of comparisons; **Figure S17**). In both cases, singlets should only be called if more than half of the methods in the combination classify the droplet as a singlet.

However, for research questions where it is crucial to remove as many doublets as possible, even if it means classifying some true singlets as doublets, the combination of *Demuxlet* and *Scds* is ideal when reference SNP genotypes are available for the individuals multiplexed in the pool (**Figure 5**). This intersectional method significantly increases the PPV compared to each method by 0.03 and an average 0.03 increase for both demultiplexing and doublet detecting methods (*t*-test FDR < 0.05). While 0.03 may appear to be a slight improvement, this change can result in 627 true singlets and doublets reclassified for a pool of 8,661 droplets - 7.2% of the total pool for *Solo* (**Figure S18**). However, this approach generally reduces the total number of true singlets classified compared to the individual methods (**Figure S18**). However, if reference SNP genotypes are not available for the individuals multiplexed in the pool, *Vireo* and *Scds* is the best combination of methods to remove as many false singlets as possible effectively - increasing the PPV by on average 0.03 compared to each of the individual methods (*t*-test FDR < 0.05), with a 0.03 average difference for both demultiplexing and double detecting methods (**Figure S19**). In both cases, singlets should only be called if more than half of the methods in the combination classify the droplet as a singlet (**Figure 5**).

#### Combining Multiple Demultiplexing Methods Improves Doublet Removal for Large Multiplexed Experimental Designs

For experiments that multiplex more than 16 individuals, we considered the combinations between *Demuxlet, Freemuxlet, Souporcell* and *Vireo* (**Table S11**) since only a small proportion of the doublets would be undetectable by demultiplexing methods (droplets that are homogenic; **Figure 3b**). To balance doublet removal and maintain true singlets, we recommend the combination of either *Demuxlet, Freemuxlet* and *Vireo* or *Demuxlet, Freemuxlet, Souporcell* and *Vireo*. These method combinations significantly increase the MCC by, on average, 0.21 compared to all the individual methods (*t*-test FDR < 0.05; **Figures S20a**). This substantially increases true singlets and true doublets relative to the individual methods (**Figures S20a**). If reference SNP genotypes are not available for the individuals multiplexed in the pools, the combination of *Freemuxlet* and *Souporcell* (16 multiplexed individuals) or *Freemuxlet, Souporcell* and *Vireo* (≥ 16 multiplexed individuals; **Figure 5**). This combinatorial approach results in a significant increase in the MCC (by 0.21 on average) compared to all the individual methods (*t*-test FDR < 0.05 for 83% of comparisons; **Figure S21** Further, for research questions where it is essential to remove as many doublets as possible, we recommend either *Demuxlet* and *Vireo* or *Demuxlet* and *Souporcell* (**Figure 5**). These combinations significantly increase 0.07 PPV on average compared to individual methods (*t*-test FDR < 0.05). Using these combinations decreases the number of true singlets classified while increasing the number of true doublets relative to individual methods (**Figures S20b**). However, if reference SNP genotypes are unavailable for the individuals multiplexed in the pool, *Freemuxlet* and *Vireo* is the best intersectional method to increase the PPV compared to the individual techniques resulting in an average improvement of 0.07 PPV (*t*-test FDR < 0.05; **Figure S21b**). Again, the best intersectional method is to call a singlet only when more than half the methods classify the droplet as a singlet (**Figure 4**).

These results collectively demonstrate that, regardless of the experimental design, demultiplexing and doublet detecting approaches that intersect multiple methods significantly enhance droplet classification. This is consistent across different pool sizes and will improve singlet annotation.

### Demuxafy Improves Doublet Removal and Improves Usability

To make our intersectional approaches accessible to other researchers, we have developed *Demuxafy* (https://demultiplexing-doublet-detecting-docs.readthedocs.io/en/latest/index.html) - an easy-to-use software platform powered by Singularity. This platform provides the requirements and instructions to execute each demultiplexing and doublet detecting methods. In addition, *Demuxafy* provides wrapper scripts that simplify method execution and effectively summarise results. We also offer tools that help expected estimate numbers of doublets and provide method combination recommendations based on scRNA-seq pool characteristics. *Demuxafy* also combines the results from multiple different methods, provides classification combination summaries, and provides final integrated combination classifications based on the intersectional techniques selected by the user. The significant advantages of *Demuxafy* include a centralised location to execute each of these methods, simplified ways to combine methods with an intersectional approach, and summary tables and figures that enable practical interpretation of multiplexed datasets (**Figure 1a**).

## Discussion

Demultiplexing and doublet detecting methods have made large-scale scRNA-seq experiments achievable. However, many demultiplexing and doublet detecting methods have been developed in the recent past, and it is unclear how their performances compare. Further, the demultiplexing techniques best detect heterogenic doublets while doublet detecting methods identify heterotypic doublets. Therefore, we hypothesised that demultiplexing and doublet detecting methods would be complementary and be more effective at removing doublets than demultiplexing methods alone.

Indeed, we demonstrated the benefit of utilising a combination of demultiplexing and doublet detecting methods. The optimal intersectional combination of methods depends on the experimental design and capture characteristics. Our results suggest super loaded captures - where a high percentage of doublets is expected - will benefit from multiplexing. Further, when many donors are multiplexed (>16), doublet detecting is not required as there are few doublets that are homogenic and heterotypic.

We have provided two different method combination recommendations based on the experimental design and whether removing doublets should be adequately balanced with maintaining a high proportion of singlets or whether it is more important to remove as many doublets as possible. This decision is highly dependent on the research question. However, we expect that the balanced approach will be appropriate for most research questions and only research questions that are interrogating extremely small effect sizes or transitional states will require the more stringent doublet removal approach. Overall, our results provide researchers with important demultiplexing and doublet detecting performance assessments and combinatorial recommendations. Our software platform, *Demuxafy* (https://demultiplexing-doublet-detecting-docs.readthedocs.io/en/latest/index.html), provides a simple implementation of our methods in any research lab around the world, providing cleaner scRNA-seq datasets and enhancing interpretation of results.

## Materials and Methods

### Data

All data have been described previously^15^. Briefly, all work was approved by the Royal Hobart Hospital, the Hobart Eye Surgeons Clinic, Human Research Ethics Committees of the Royal Victorian Eye and Ear Hospital (11/1031), University of Melbourne (1545394) and University of Tasmania (H0014124) in accordance with the requirements of the National Health & Medical Research Council of Australia (NHMRC) and conformed with the Declaration of Helsinki^17^.

### PBMC scRNA-seq Data

Blood samples were collected and processed as described previously^18^. Briefly, mononuclear cells were isolated from whole blood samples and stored in liquid nitrogen until thawed for scRNA-seq capture. Equal numbers of cells from 12-16 samples were multiplexed per pool and single-cell suspensions were super loaded on a Chromium Single Cell Chip A (10x Genomics) to capture 20,000 droplets per pool. Single-cell libraries were processed per manufacturer instructions and the 10x Genomics Cell Ranger Single Cell Software Suite (v 2.2.0) was used to process the data and map it to GRCh38. The quality control metrics of each pool are demonstrated in **Figure S22**.

### PBMC DNA SNP Genotyping

SNP genotype data were prepared as described previously^18^. Briefly, DNA was extracted from blood with the QIAamp Blood Mini kit and genotyped on the Illumina Infinium Global Screening Array. SNP genotypes were processed with Plink and GCTA before imputing on the Michigan Imputation Server using Eagle v2.3 for phasing and Minimac3 for imputation based on the Haplotype Reference Consortium panel (HRCr1.1). SNP genotypes were then lifted to hg38 and filtered for > 1% minor allele frequency (MAF) and an R^2^ > 0.3.

### Fibroblast scRNA-seq Data

The fibroblast scRNA-seq data has been described previously^15^. Briefly, human skin punch biopsies from donors over the age of 18 were cultured in DMEM high glucose supplemented with 10% fetal bovine serum (FBS), L-gluatmine, 100 U/mL penicillin and 100 μg/mL (Thermo Fisher Scientific, USA).

For scRNA-seq, viable cells were flow sorted and single cell suspensions were loaded onto a 10x Genomics Single Cell 3’ Chip and were processed per 10x instructions and the Cell Ranger Single Cell Software Suite from 10x Genomics was used to process the sequencing data into transcript count tables as previously described^15^. The quality control metrics of each pool are demonstrated in **Figure S23**.

### Fibroblast DNA SNP Genotyping

The DNA SNP genotyping for fibroblast samples has been described previously^15^. Briefly, DNA from each donor was genotyped on an Infinium HumanCore-24 v1.1 BeadChip (Illumina). GenomeStudioTM V2.0 (Illumina), Plink and GenomeStudio were used to process the SNP genotypes. Eagle V2.3.5 was used to phase the SNPs and it was imputed with the Michigan Imputation server using minimac3 and the 1000 genome phase 3 reference panel as described previously^15^.

#### Demultiplexing Methods

All the demultiplexing methods were built and run from a singularity image.

### Popscle

The Popscle v0.1-beta suite^14^ for population genomics in single cell data was used for *Demuxlet* and *Freemuxlet* demultiplexing methods. The *popscle dsc-pileup* function was used to create a pileup of variant calls at known genomic locations from aligned sequence reads in each droplet with default arguments.

#### Demuxlet

*Demuxlet*^*2*^ is a SNP genotype reference-based single cell demultiplexing method. *Demuxlet* was run with a genotype error coefficient of 1 and genotype error offset rate of 0.05 and the other default parameters using the *popscle demuxlet* command from Popscle (v0.1-beta).

#### Freemuxlet

*Freemuxlet*^*14*^ is a SNP genotype reference-free single cell demultiplexing method. *Freemuxlet* was run with default parameters including the number of samples included in the pool using the *popscle freemuxlet* command from Popscle (v0.1-beta).

### ScSplit

*ScSplit* v1.0.7^2^ was downloaded from the *ScSplit* github and the recommended steps for data filtering quality control prior to running *ScSplit* were followed. Briefly, reads that had read quality lower than 10, were unmapped, were secondary alignments, did not pass filters, were optical PCR duplicates, were secondary alignments or were duplicate reads were removed. The resulting bam file was then sorted and indexed followed by freebayes to identify single nucleotide variants (SNVs) in the dataset. The resulting SNVs were filtered for quality scores greater than 30 and for variants present in the reference SNP genotype vcf. The resulting filtered bam and vcf files were used as input for the s*cSplit count* command with default settings to count the number of reference and alternative alleles in each droplet. Next the allele matrices were used to demultiplex the pool and assign cells to different clusters using the *scSplit run* command including the number of individuals (*-n*) option and all other options set to default. Finally, the individual genotypes were predicted for each cluster using the *scSplit genotype* command with default parameters.

### Souporcell

*Souporcell*^*5*^ is a SNP genotype reference-free single cell demultiplexing method. The *Souporcell* v1.0 singularity image was downloaded via instructions from the gihtub page. The *Souporcell* pipeline was run using the *souporcell_pipeline*.*py* script with default options and the option to include known variant locations (*--common_variants*).

### Vireo

*Vireo*^*3*^ is a single cell demultiplexing method that can be used with reference SNP genotypes or without them. For this assessment, *Vireo* was used with reference SNP genotypes. Per *Vireo* recommendations, we used Model 1 of the cellSNP version 0.3.2 to make a pileup of SNPs for each droplet with the recommended options using the genotyped reference genotype file as the list of common known SNP and filtered with SNP locations that were covered by at least 20 UMIs and had at least 10% minor allele frequency across all droplets. *Vireo* version 0.4.2 was then used to demultiplex using reference SNP genotypes and indicating the number of individuals in the pools.

#### Doublet Detecting Methods

All doublet detecting methods were built and run from a Singularity image.

### DoubletDecon

*DoubletDecon*^*9*^ is a transcription-based deconvolution method for identifying doublets. *DoubletDecon* version 1.1.6 analysis was run in R version 3.6.3. *SCTransform* ^19^ from Seurat ^20^ version 3.2.2 was used to preprocess the scRNA-seq data and then the *Improved_Seurat_Pre_Process* function was used to process the SCTransformed scRNA-seq data. Clusters were identified using Seurat function *FindClusters* with resolution 0.2 and 30 principal components (PCs). Then the *Main_Doublet_Decon* function was used to deconvolute doublets from singlets for six different rhops - 0.6, 0.7, 0.8, 0.9, 1.0 and 1.1. Then the rhop that resulted in the closest number of doublets to the expected number of doublets per the following equation:

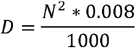

where *N* is the number of droplets captured and *D* is the number of expected doublets.

### DoubletDetection

*DoubletDetection*^*7*^ is a transcription-based method for identifying doublets. *DoubletDetection* version 2.5.2 analysis was run in python version 3.6.8. Droplets without any UMIs were removed before analysis with *DoubletDetection*. Then the *doubletdetection*.*BoostClassifier* function was run with 50 iterations with *use_phenograph* set to False and *standard_scaling* set to True. The predicted number of doublets per iteration was visualized across all iterations and any pool that did not converge after 50 iterations, it was run again with increasing numbers of iterations until they reached convergence.

### DoubletFinder

*DoubletFinder*^*8*^ is a transcription-based doublet detecting method. *DoubletFinder* version 2.0.3 was implemented in R version 3.6.3. First, droplets that were more than 3 median absolute deviations (mad) away from the median for mitochondrial per cent, ribosomal per cent, number of UMIs or number of genes were removed per developer recommendations. Then the data was normalized with SCTransform followed by cluster identification using *FindClusters* with resolution 0.3 and 30 principal components (PCs). Then, pKs were selected by the pK that resulted in the largest BC_MVN_. Finally, the homotypic doublet proportions were calculated and the number of expected doublets with the highest doublet proportion were classified as doublets per the following equation:

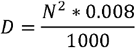

where N is the number of droplets captured and D is the number of expected doublets.

### ScDblFinder

*ScDblFinder*^*10*^ is a transcription-based method for detecting doublets from scRNA-seq data. *ScDblFinder* 1.3.25 was implemented in R version 4.0.3. *ScDblFinder* was implemented with two sets of options. The first included implementation with the expected doublet rate as calculated by:

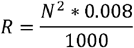

where *N* is the number of droplets captured and *R* is the expected doublet rate. The second condition included the same expected number of doublets and included the doublets that had already been identified by all the demultiplexing methods.

### Scds

*Scds*^*11*^ is a transcription-based doublet detecting method. Scds version 1.1.2 analysis was completed in R version 3.6.3. *Scds* was implemented with the *cxds* function and *bcds* functions with default options followed by the *cxds_bcds_hybrid* with *estNdbl* set to TRUE so that doublets will be estimated based on the values from the *cxds* and *bcds* functions.

### Scrublet

*Scrublet*^*12*^ is a transcription-based doublet detecting method for single-cell RNA-seq data. *Scrublet* was implemented in python version 3.6.3. *Scrublet* was implemented per developer recommendations with at least three counts per droplet, three cells expressing a given gene, 30 PCs and a doublet rate based on the following equation:

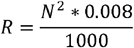

where *N* is the number of droplets captured and *R* is the expected doublet rate. Four different minimum number of variable gene percentiles: 80, 85, 90 and 95. Then, the best variable gene percentile was selected based on the distribution of the simulated doublet scores and the location of the doublet threshold selection. In the case that the selected threshold does not fall between a bimodal distribution, those pools were run again with a manual threshold set.

### Solo

*Solo*^*13*^ is a transcription-based method for detecting doublets in scRNA-seq data. *Solo* was implemented with default parameters and an expected number of doublets based on the following equation:

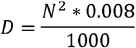

where N is the number of droplets captured and D is the number of expected doublets. *Solo* was additionally implemented in a second run for each pool with the doublets that were identified by all the demultiplexing methods as known doublets to initialize the model.

#### In Silico Pool Generation

Cells that were identified as singlets by all methods were used to simulate pools with the *synth_pool*.*py* script provided by the Vireo ^3^ package. Ambient RNA was simulated by changing the barcodes and UMIs on a random selection of reads for 2, 5 or 10% of the total UMIs. High mitochondrial per cent simulations were produced by replacing reads in 5, 10 or 25% of the randomly selected cells with mitochondrial reads. The number of reads to replace was derived from a normal distribution with an average of 30 and a standard deviation of three. Decreased read coverage of pools was simulated by down-sampling the reads by two-thirds of the original coverage.

#### Classification Annotation

### Demultiplexing Methods

Demultiplexing methods classifications were considered correct if the droplet annotation (singlet or doublet) and the individual annotation was correct. If the droplet type was correct but the individual annotation was incorrect (*i*.*e*., classified as a singlet but annotated as the wrong individual), then the droplet was incorrectly classified.

### Doublet Detecting Methods

Doublet detecting methods were considered to have correct classifications if the droplet annotation matched the known droplet type.

#### Analyses

All downstream analyses were completed in R version 4.0.2.

