## Supplementary figures and text for "*Demuxafy*: Improvement in droplet assignment by integrating multiple single-cell demultiplexing and doublet detection methods"

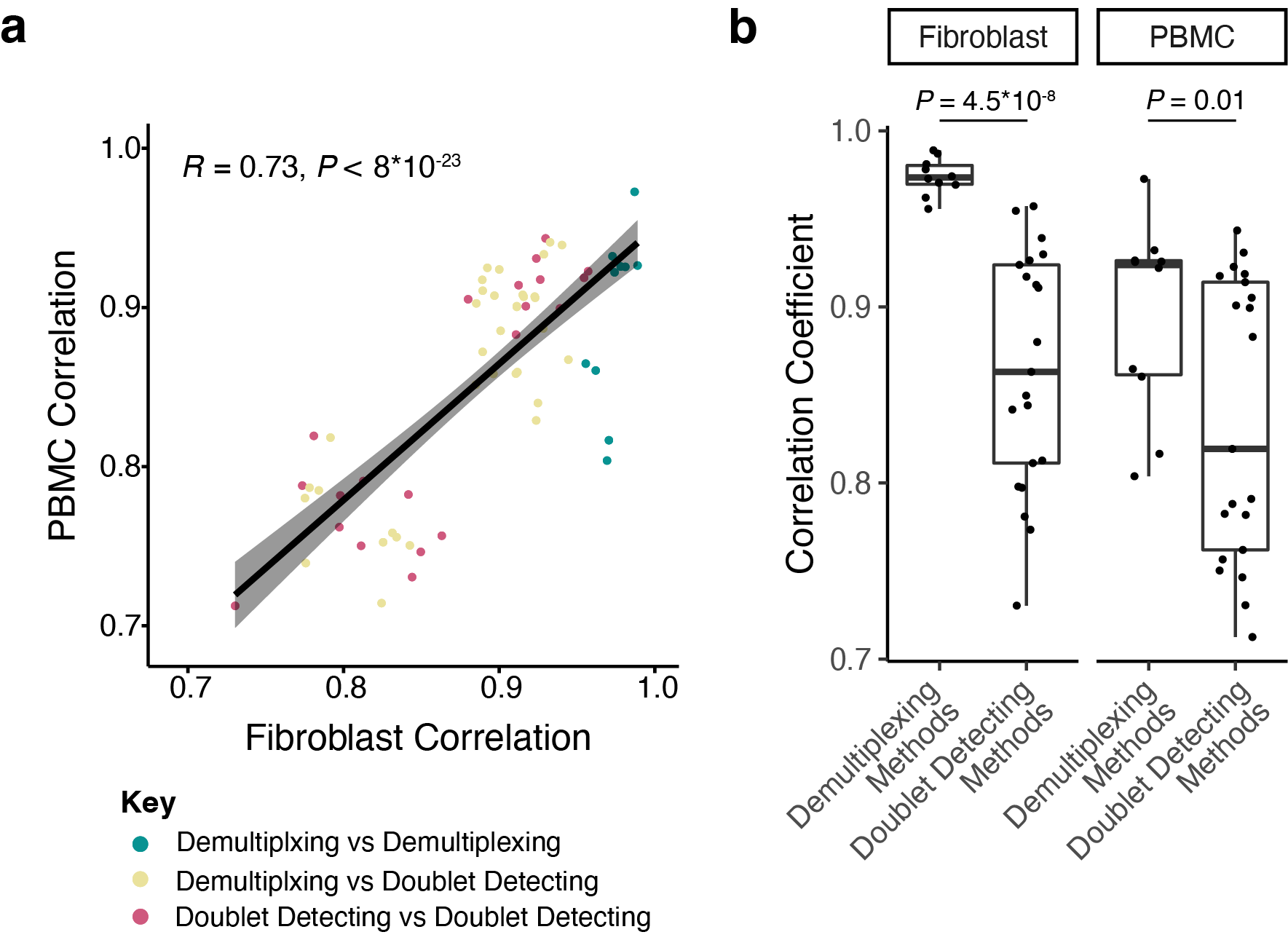

**Supplementary Figure 1:** **Association of Method Correlations Between PBMC and Fibroblast Cell Types**. **a**) The pairwise correlations between the demultiplexing and doublet detecting methods were tested for association between the PBMC and fibroblast cell types. The colours indicate which method types are being compared in the correlation. Spearman correlation was used to test the relationship between the PBMC and Fibroblast correlations. **b**) Comparison of the correlations between the demultiplexing and doublet detecting methods. Distributions were tested with a one-sided Wilcoxon rank-sum test.

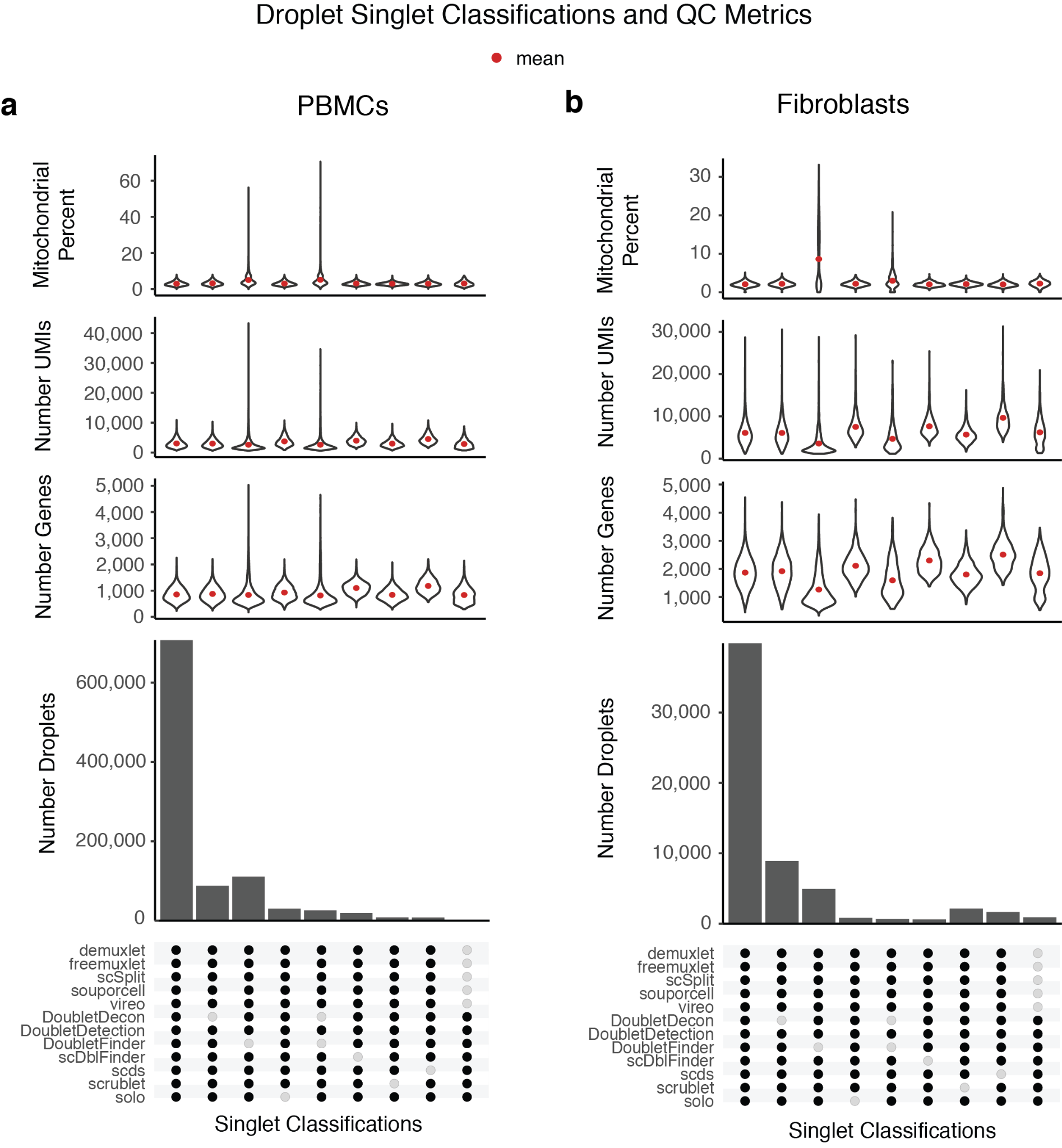

**Supplementary Figure 2: Cell Type Metrics for Droplets Classified as Singlets by Different Method Combinations.** The distribution of mitochondrial per cent, number of UMIs and number of genes and demonstrated with violin plots for the top method classification combinations in PBMCs (**a**) and fibroblasts (**b**).

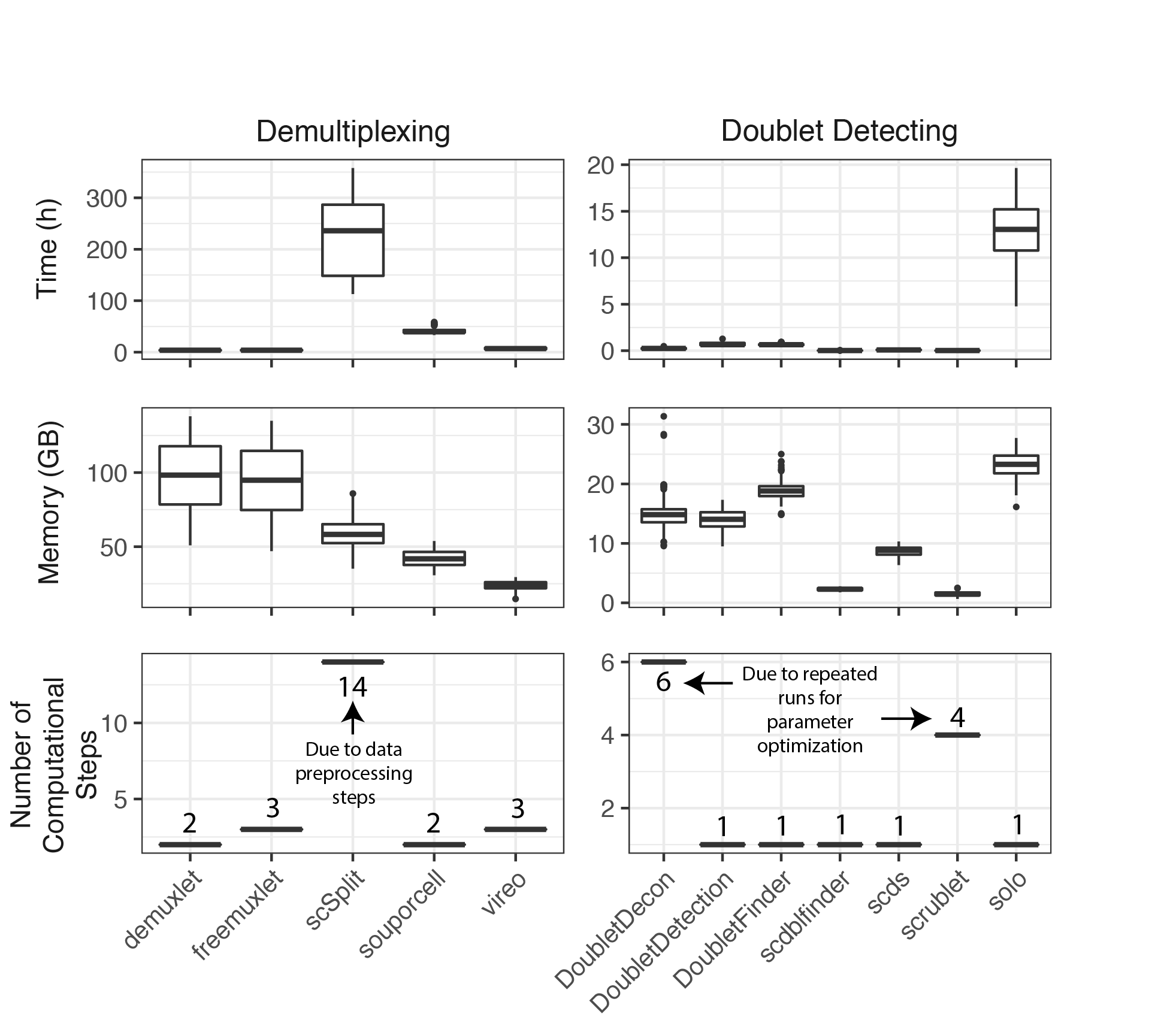

**Supplementary Figure 3: The time, memory and number of computational steps for each method.** The time in hours (h), the memory in gigabytes (GB) and the number of user-required computational steps for each of the demultiplexing (left) and doublet detecting methods (right).

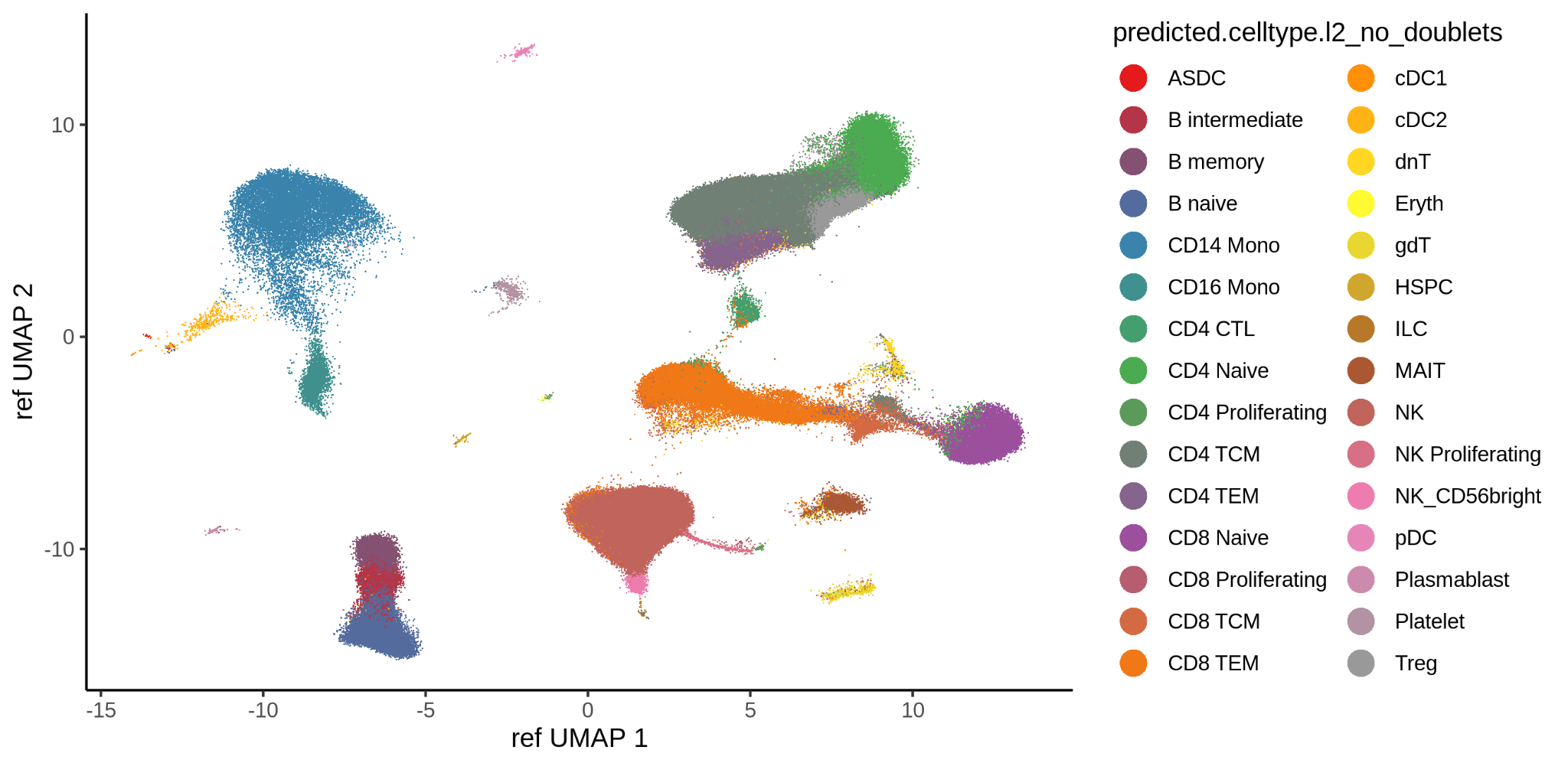

**Supplementary Figure 4: Cell Type Annotation of droplets classified as singlets by all methods.** The cell type annotation of the PBMCs that were all classified as singlets by all the demultiplexing and doublet detecting methods using the Azimuth PBMC reference.

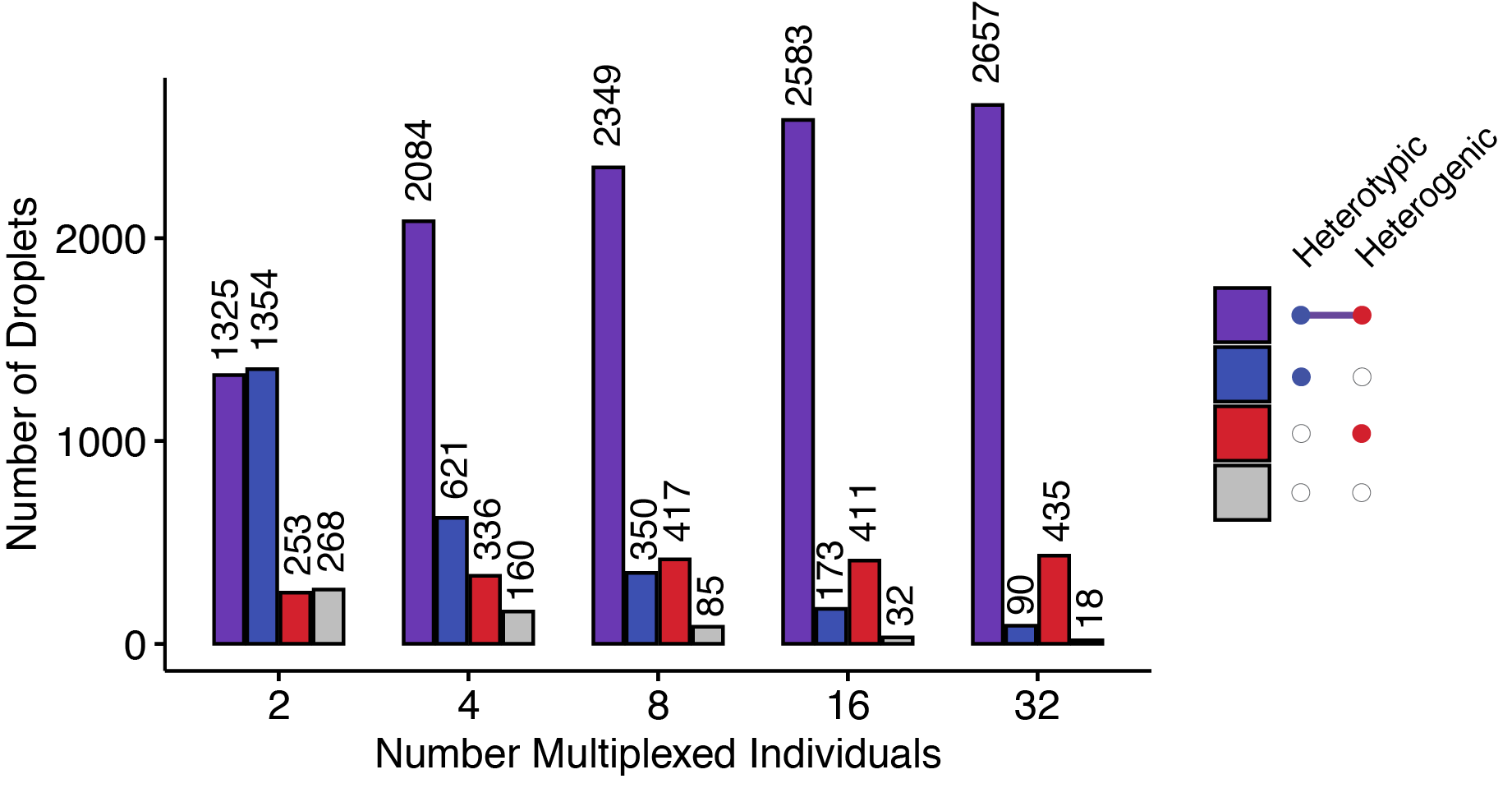

**Supplementary Figure 5: Number of different doublet types assuming 20,000 droplets captured.**  The number of different types of doublets would be in pools that multiplexed 2, 4, 8, 16 and 32, assuming 20,000 droplets were captured.

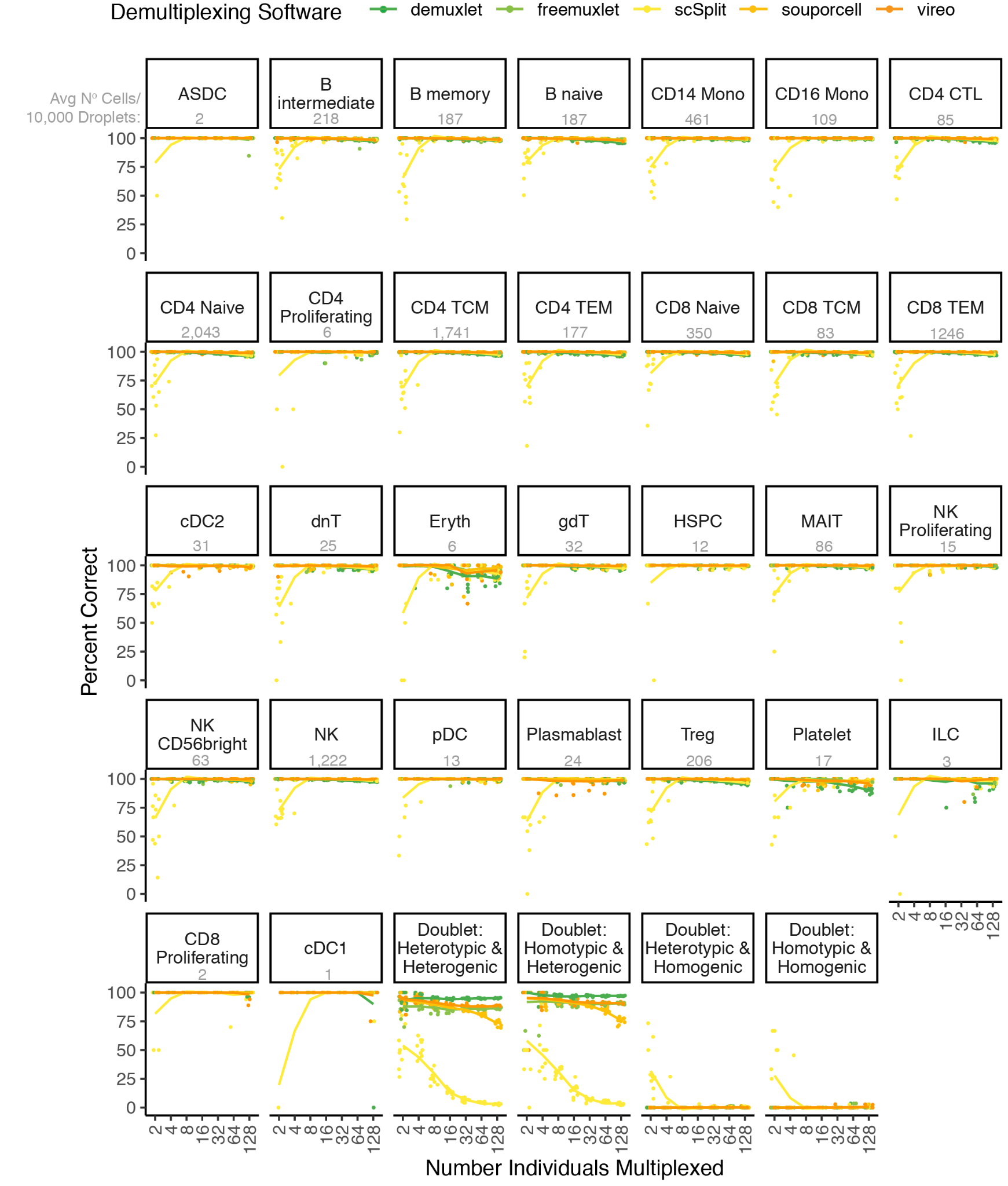

**Supplementary Figure 6: Each demultiplexing method's percentage of each cell type annotated correctly—the** percentage of droplets correctly classified by each demultiplexing method across different multiplexed pool sizes. The number of each cell type per 10,000 droplets is shown in grey beneath the cell type name. N = 10 for each group.

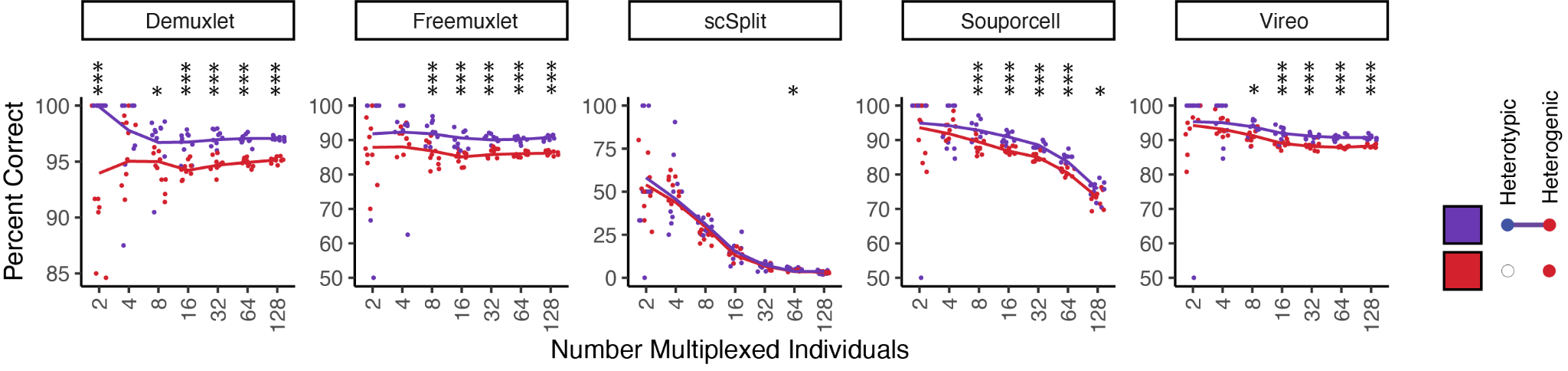

**Supplementary Figure 7: Difference in per cent correct for heterogenic doublets that are heterotypic and homotypic with Demultiplexing Methods.** Most of the demultiplexing methods (except *ScSplit*) are significantly more effective at detecting heterogenic doublets that are also heterotypic (purple) as opposed to homotypic (red). N = 10 for each group. Statistical significance was tested with a student’s *t*-test for each multiplexed pool size for each method. *** *P* < 0.001, ** 0.001 < *P* < 0.01, * 0.01 < *P* < 0.05.

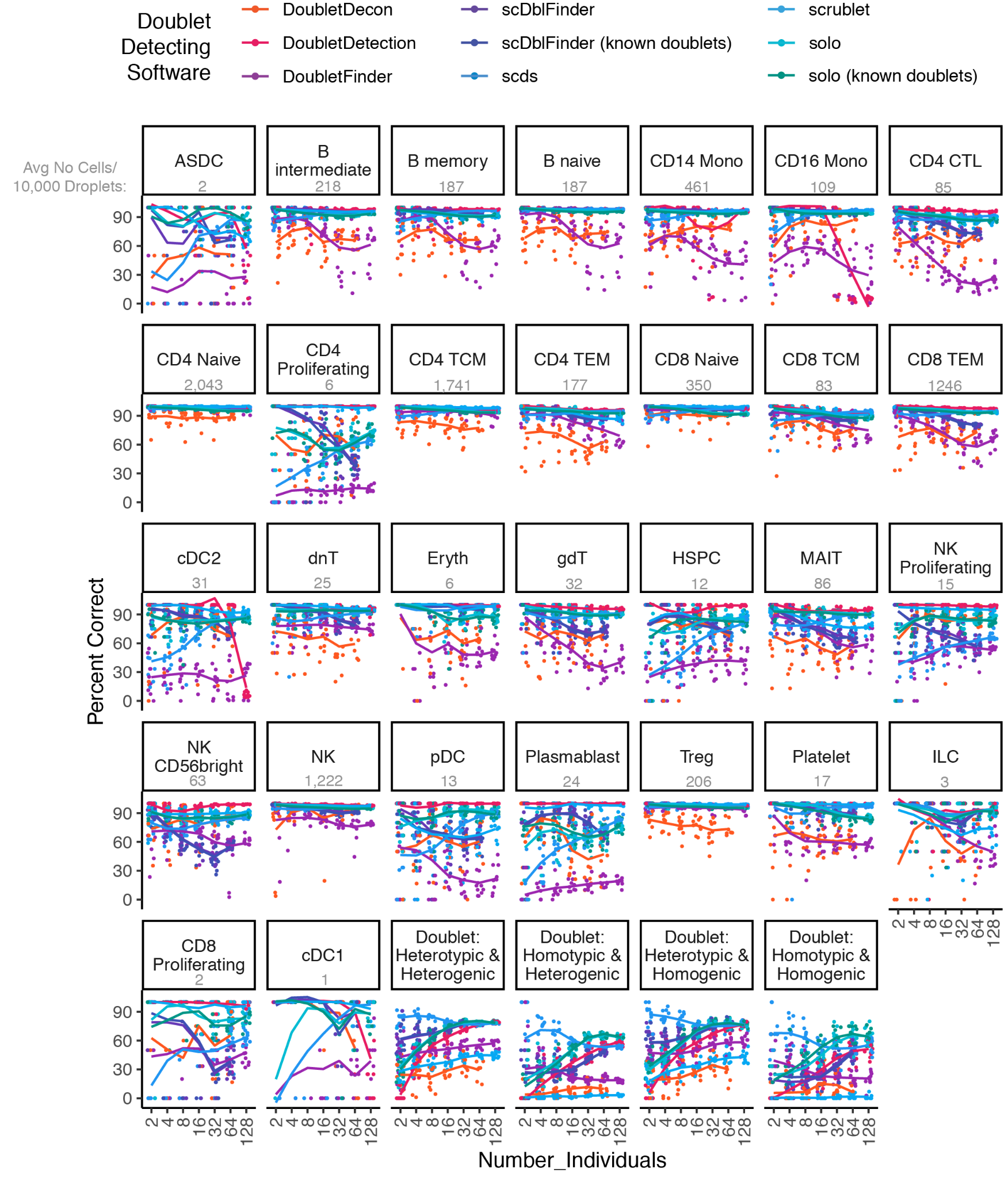

**Supplementary Figure 8: The percentage of each doublet detecting method’s cell type annotated correctly—the** percentage of droplets correctly classified by each doublet detecting method across different multiplexed pool sizes. The number of each cell type per 10,000 droplets is shown in grey beneath the name of each cell type. N = 10 for each group.

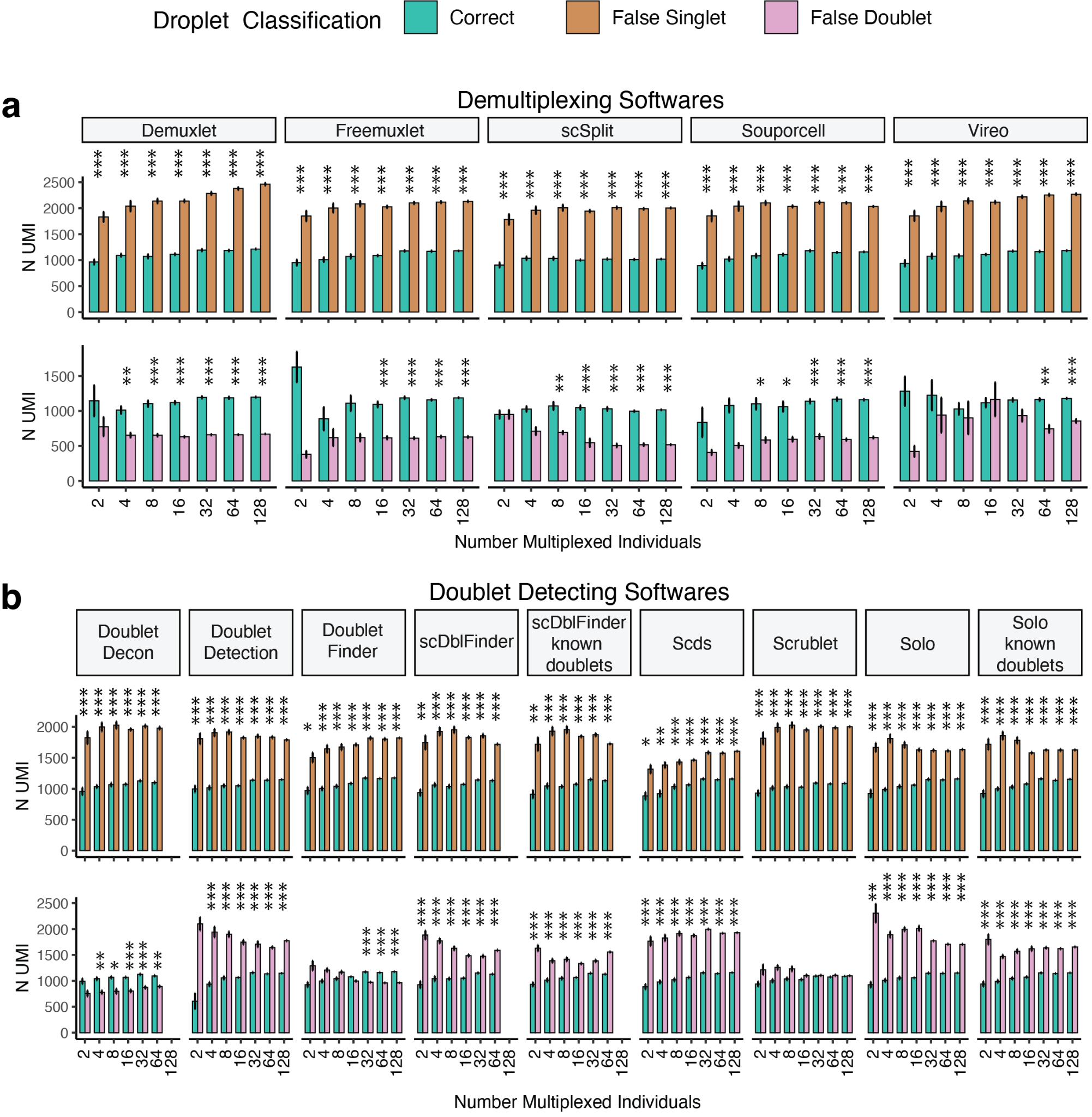
**Supplementary Figure 9: Number of UMIs for False Doublets and Singlets Compared to Correctly Classified Droplets.** **a**) The false singlet droplets had higher UMI counts than the correctly identified droplets for all demultiplexing methods for all pool sizes. The false doublets demonstrated lower UMI counts for most methods and pool sizes. The exceptions were smaller pools and Vireo, which only demonstrated significant differences for pools that multiplexed two, 32, 64 and 128 individuals. **b**) Similar to the demultiplexing methods, the falsely identified singlets by doublet detecting methods had higher UMI counts than the correctly classified droplets. The false singlets identified by *DoubletDetection*, *scDblFinder*, *scDblFinder* with known doublets, *Solo* and *Solo* with known doublets had higher UMI counts than the correct droplets. The false singlets identified by *DoubletFinder* and *Scrublet* had higher UMI counts for smaller pools, and those identified by *DoubletDecon* and large pools with *DoubletFinder* had significantly lower UMIs than the correctly identified droplets. The difference in the number of UMIs between correct and incorrect droplet annotation was tested with a student’s *t*-test, and *P*-values were corrected for multiple testing with the Bonferroni method. * 0.01 < P < 0.05; ** 0.001 < P < 0.01; *** P < 0.0001.

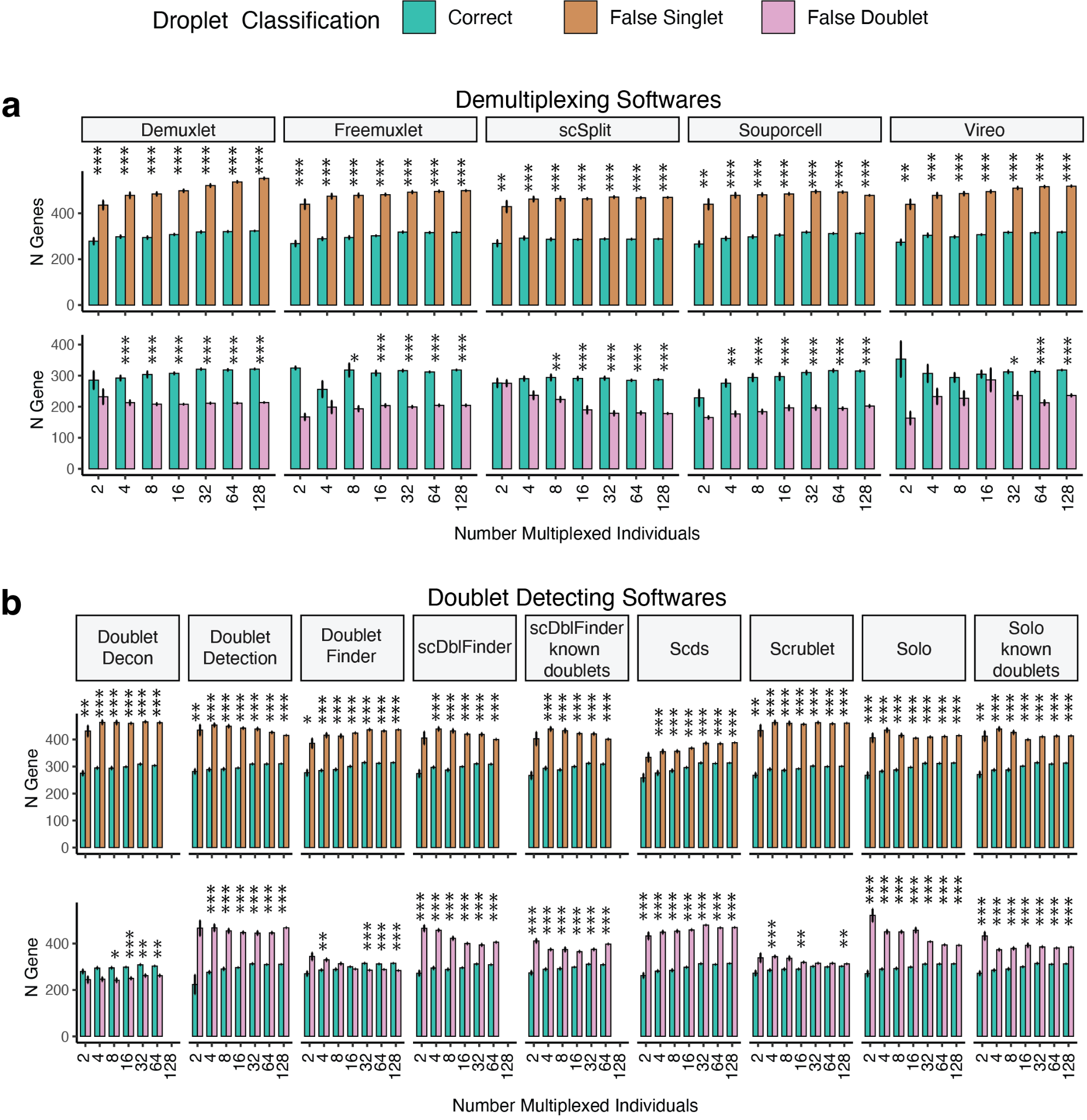

**Supplementary Figure 10: Number of Genes in False Doublets and Singlets Compared to Correctly Classified Droplets.** **a**) The false singlet droplets had higher gene counts than the correctly identified droplets for all demultiplexing methods for all pool sizes. The false doublets demonstrated lower gene counts for most methods and pool sizes. The exceptions were smaller pools and Vireo, which only demonstrated significant differences for pools that multiplexed eight, 32, 64 and 128 individuals. **b**) Like the demultiplexing methods, the falsely identified singlets by doublet detecting methods had higher gene counts than the correctly classified droplets. The false singlets identified by *DoubletDetection*, *scDblFinder*, *scDblFinder* with known doublets, *Scrublet*, *Solo* and *Solo* with known doublets all had higher gene counts than the correct droplets. The false singlets identified by *DoubletFinder* had higher gene counts for smaller pools, and those identified by *DoubletDecon* and large pools with *DoubletFinder* had significantly fewer genes than the correctly identified droplets. The difference in genes between correct and incorrect droplet annotation was tested with a student’s t-test, and P-values were corrected for multiple testing using the Bonferroni method. * 0.01 < P < 0.05; ** 0.001 < P < 0.01; *** P < 0.0001.

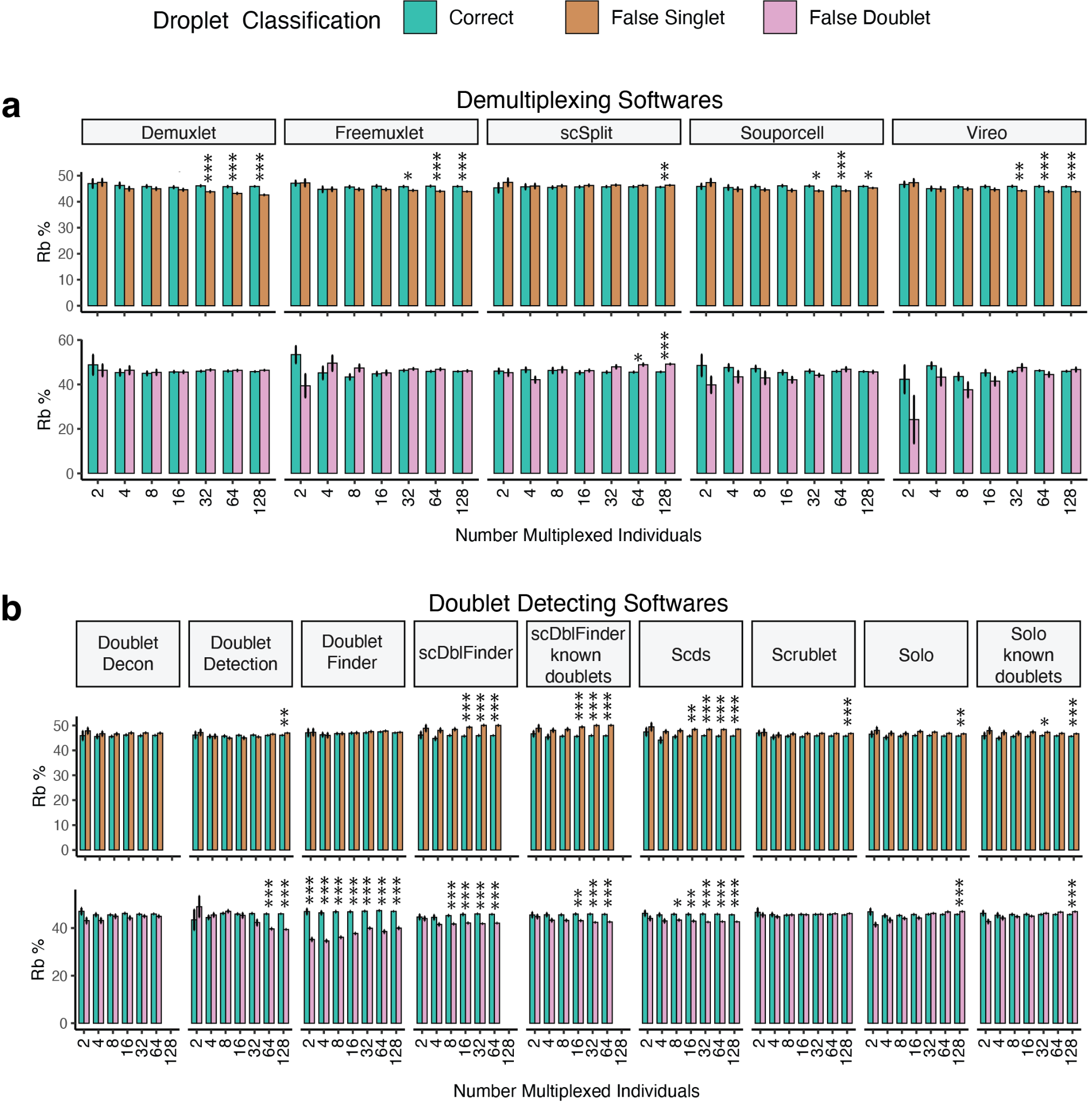

**Supplementary Figure 11: Ribosomal Percent in False Doublets and Singlets Compared to Correctly Classified Droplets.** **a**) The ribosomal per cent in false singlet droplets were consistent with the correctly identified droplets for the demultiplexing methods except for larger pool sizes which demonstrated small but significant differences in ribosomal per cent. The false doublets demonstrated consistent ribosomal per cent except for a few pool sizes annotated by *Demuxlet* and *scSplit,* which demonstrated small but significant differences. **b**) The doublet detecting methods demonstrated small but significantly higher ribosomal percent in the false doublets than the correctly classified droplets - the opposite direction of the demultiplexing methods. However, this effect was not as noticeable in *DoubletDecon*, *DoubletDetection* and *DoubletFinder*. The false doublets were less consistent across different doublet detecting methods. Some methods demonstrated significantly lower ribosomal percent in the false doublets (*i.e.* *DoubletDetection*, *DoubletFinder*, SciFinder and *scDblFinder* with known doublets). In contrast, other methods demonstrated significantly higher ribosomal percent in the falsely identified doublets (*i.e.* *Solo* and *Solo* with known doublets). The difference in genes between correct and incorrect droplet annotation was tested with a student’s t-test, and P-values were corrected for multiple testing using the Bonferroni method. * 0.01 < P < 0.05; ** 0.001 < P < 0.01; *** P < 0.0001.

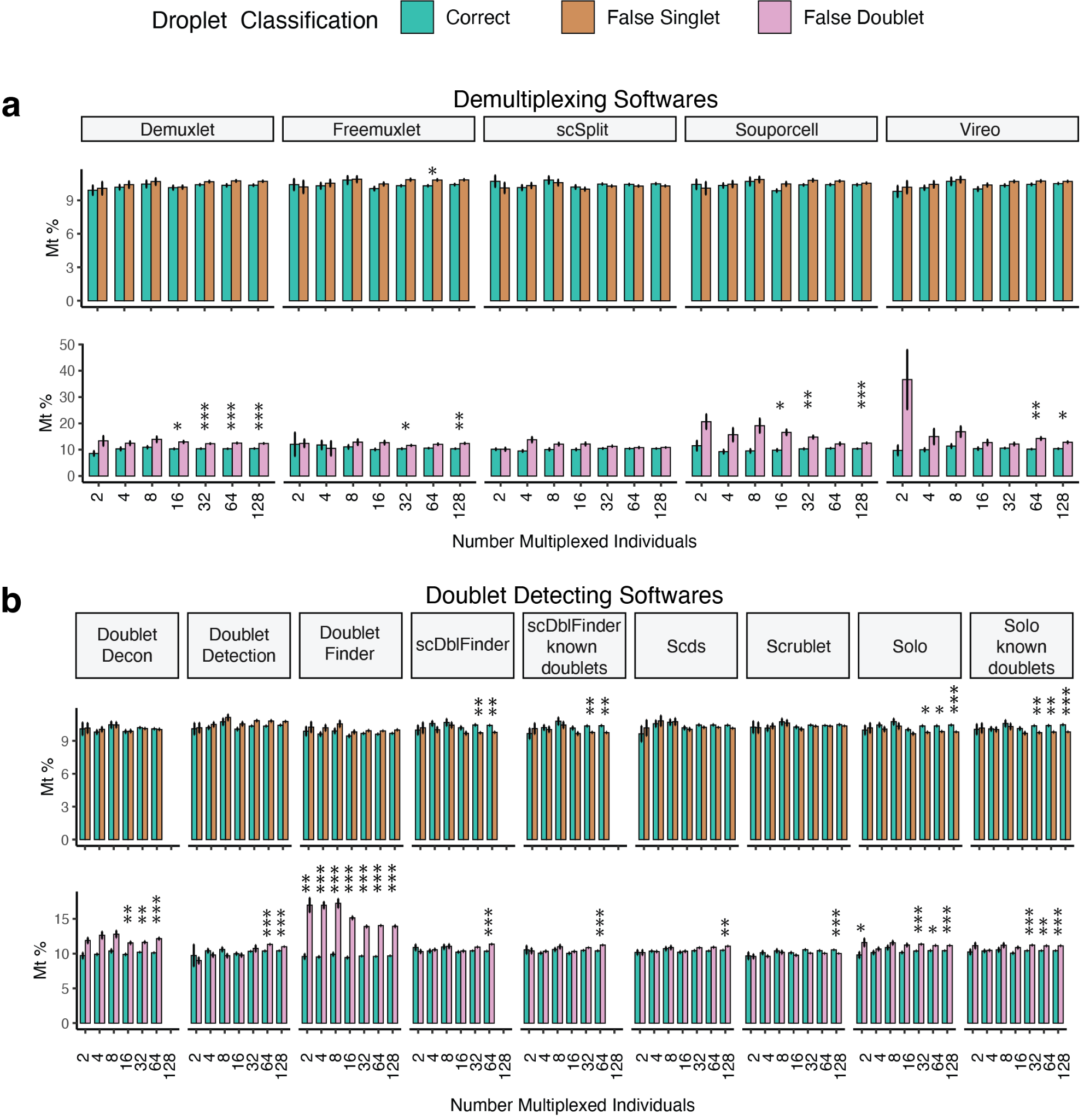

**Supplementary Figure 12: Mitochondrial Percent in False Doublets and Singlets Compared to Correctly Classified Droplets.** **a**) Only a few of the demultiplexing methods for a subset of pool sizes demonstrated significantly higher mitochondrial percent in the false singlets than the correctly classified droplets. However, the false doublets had a higher mitochondrial percent for many pool sizes across all demultiplexing methods. **b**) The doublet detecting methods demonstrated small but significantly lower mitochondrial percent for large pool sizes for false doublets - the opposite direction of the demultiplexing methods. However, this effect was not as noticeable in *DoubletDecon* or *Scrublet*. The false doublets identified by the doublet detecting methods had higher mitochondrial percent than the correctly classified droplets (especially for larger pool sizes) except for *Scrublet,* whose false doublets had lower mitochondrial percent. The difference in genes between correct and incorrect droplet annotation was tested with a student’s t-test, and P-values were corrected for multiple testing using the Bonferroni method. * 0.01 < P < 0.05; ** 0.001 < P < 0.01; *** P < 0.0001.

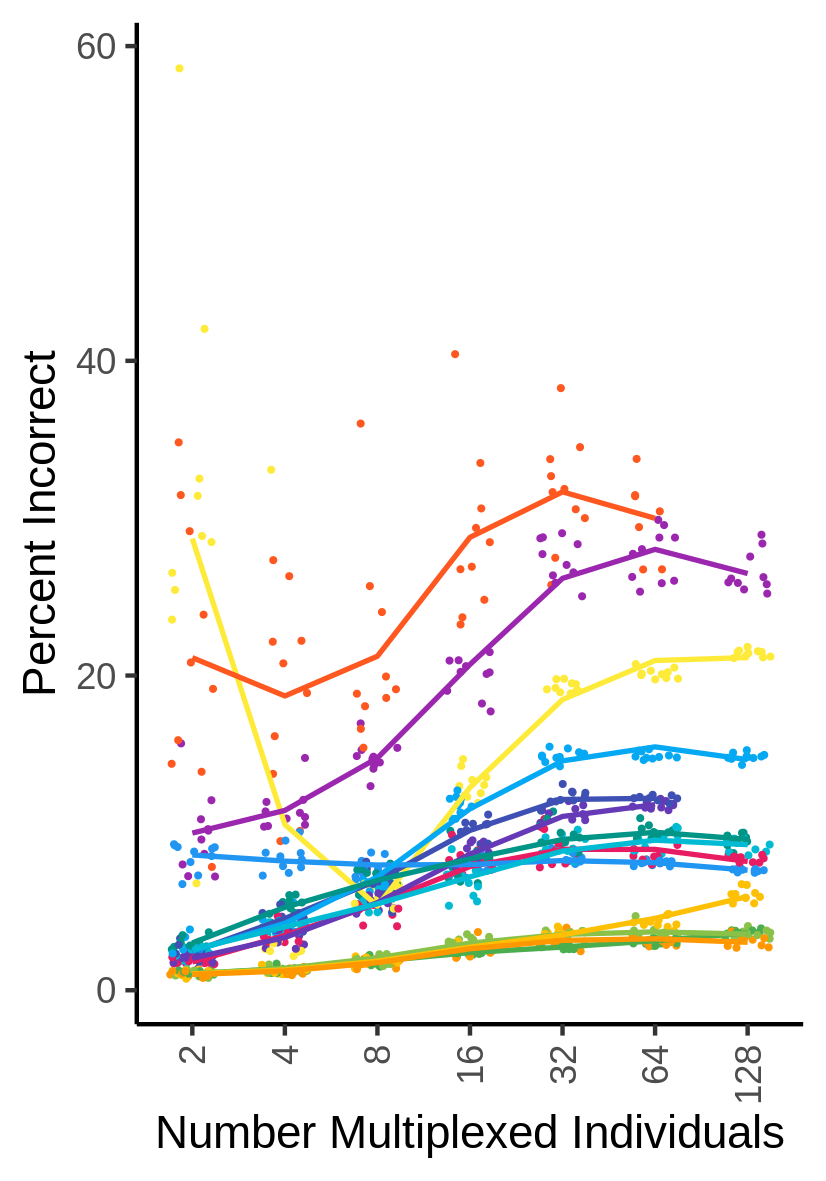

**Supplementary Figure 13: Percent Incorrect Droplet Classifications.** The percent of droplets incorrectly classified for each method across multiple multiplexed pool sizes. For demultiplexing methods, the droplet type and individual assignment had to be correct for the overall assignment to be considered valid.

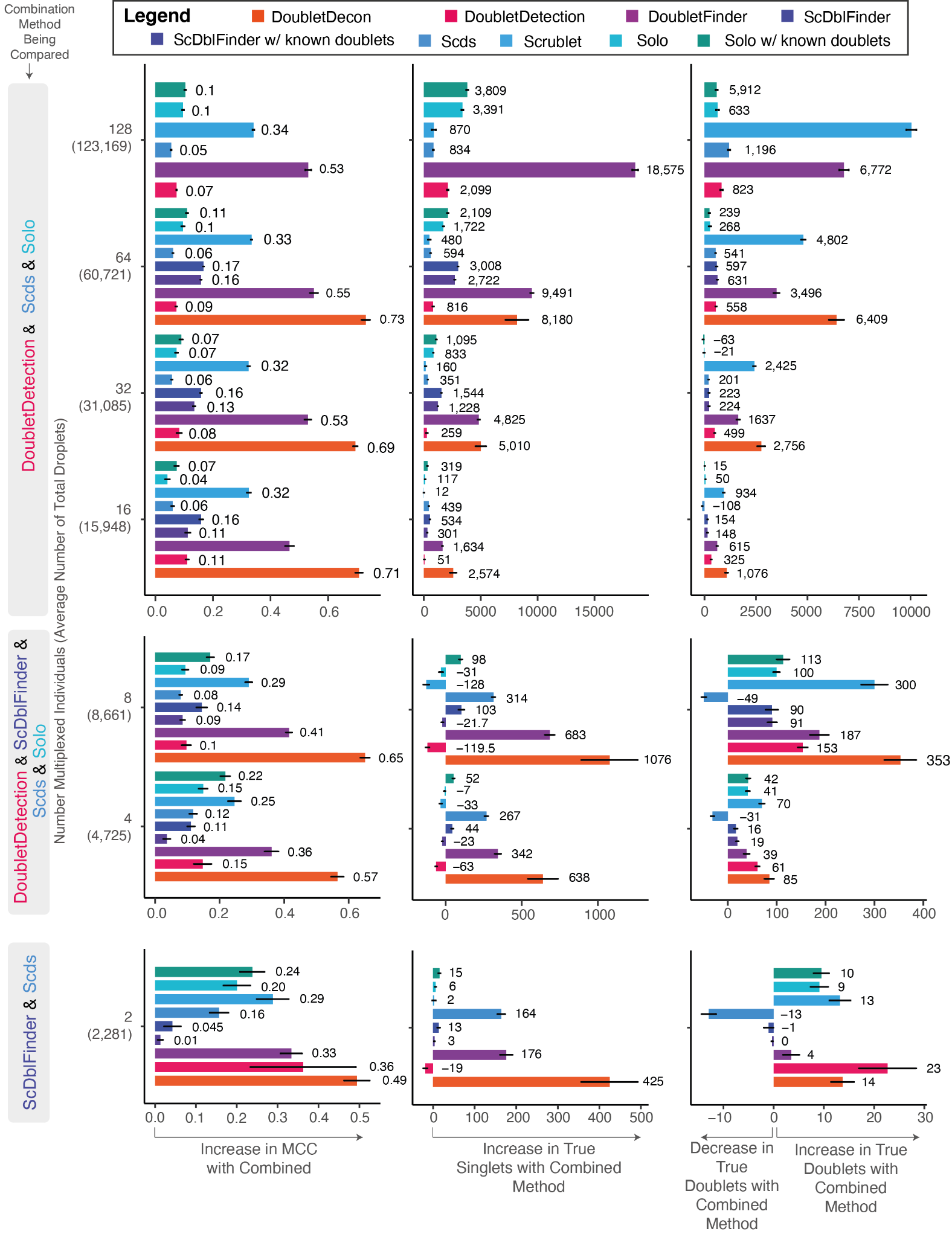

**Supplementary Figure 14: Change in MCC, True Singlets and True Doublets with Intersectional Methods Compared to Individual Methods for Non-multiplexed Pools.**
The average increase in the MCC (left) and the resulting changes in true singlets (middle) and true doublets (right) when the recommended intersectional methods for non-multiplexed pools compared to each of the individual software when aiming to use a balanced approach to remove true doublets but not at the expense of eliminating too many true singlets. Numbers are represented as averages of method performance across ten pools for each multiplexed pool number. Error bars represent the standard error around the average across the ten pools for each pool size.

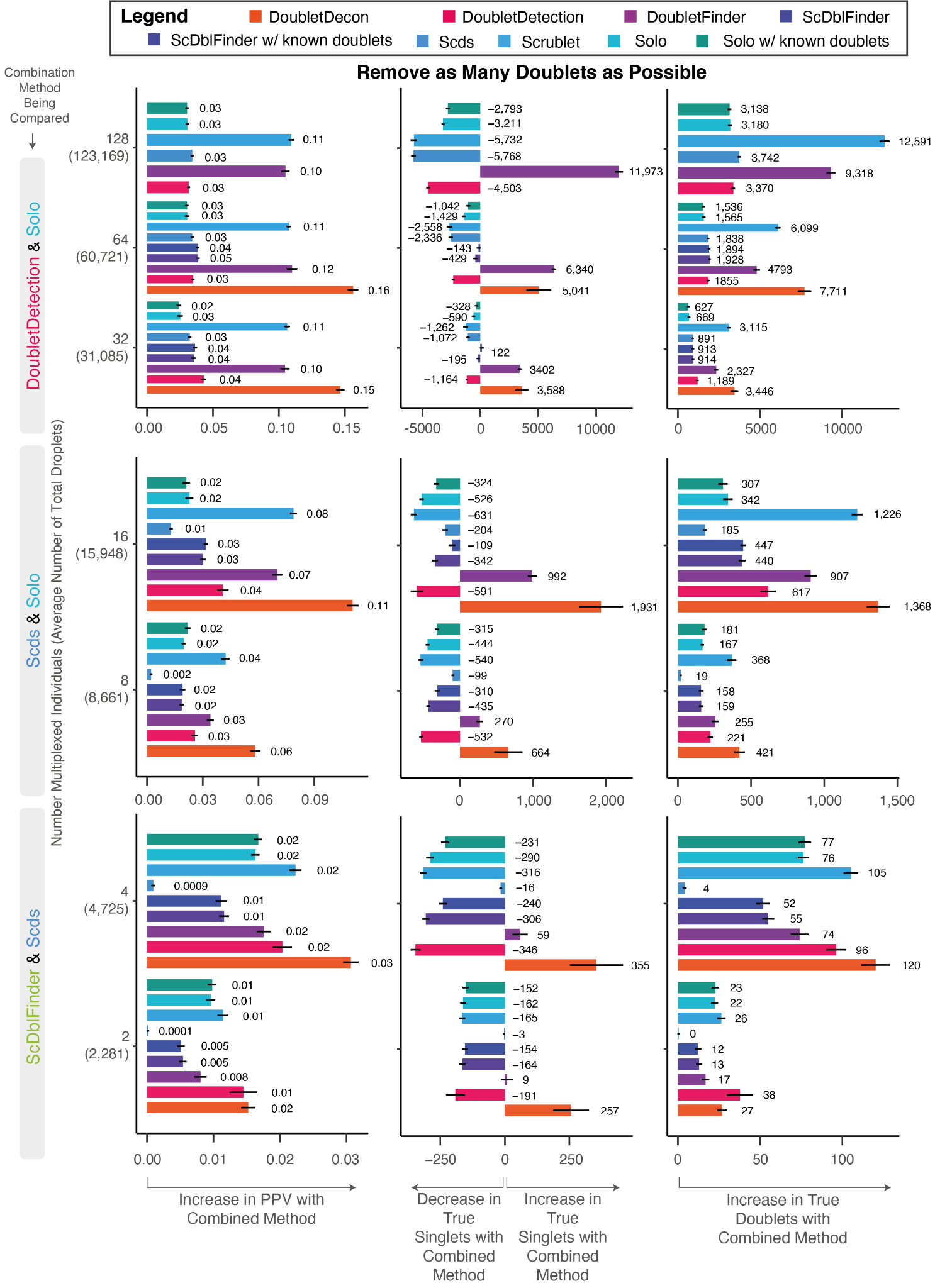

**Supplementary Figure 15: Change in PPV, True Singlets and True Doublets with Intersectional Methods Compared to Individual Methods for Non-multiplexed Pools. The average** increase in the PPV (left) and the resulting changes in true singlets (middle) and true doublets (right) when the recommended intersectional methods for non-multiplexed pools compared to each of the individual software when aiming to remove as many false singlets as possible. Numbers are represented as averages of method performance across ten pools for each multiplexed pool number. Error bars represent the standard error around the average across the ten pools for each pool size.

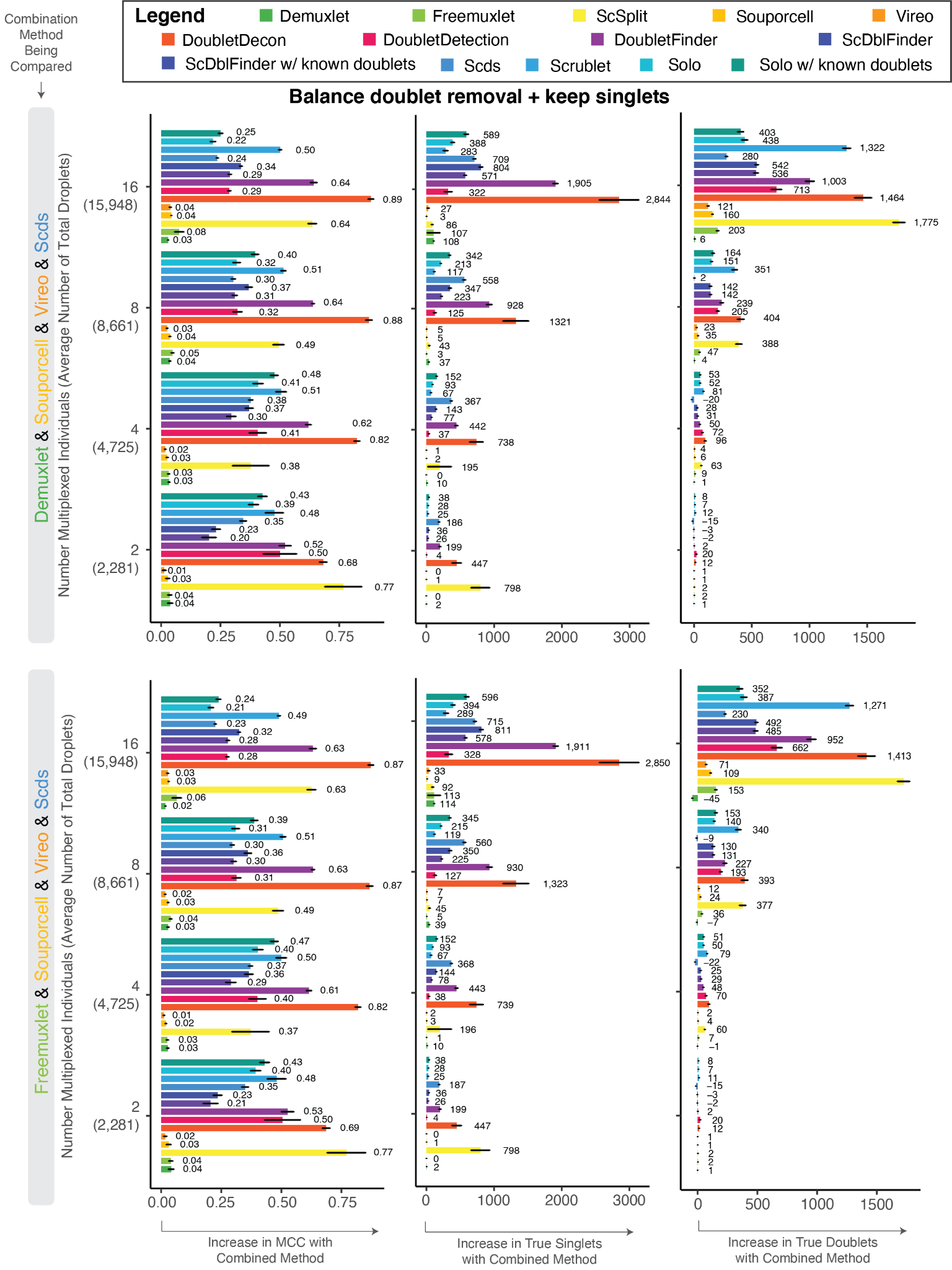

**Supplementary Figure 16: Change in MCC, True Singlets and True Doublets with Intersectional Methods Compared to Individual Methods for Multiplexed Pools with 16 or Fewer Individuals. The average** increase in the MCC (left) and the resulting changes in true singlets (middle) and true doublets (right) for the recommended intersectional methods compared to the individual techniques for multiplexed pools with 16 or fewer individuals when aiming to use a balanced approach to remove true doublets but not at the expense of eliminating too many true singlets. Numbers are represented as averages of method performance across ten pools for each multiplexed pool number. Error bars represent the standard error around the average across the ten pools for each pool size.

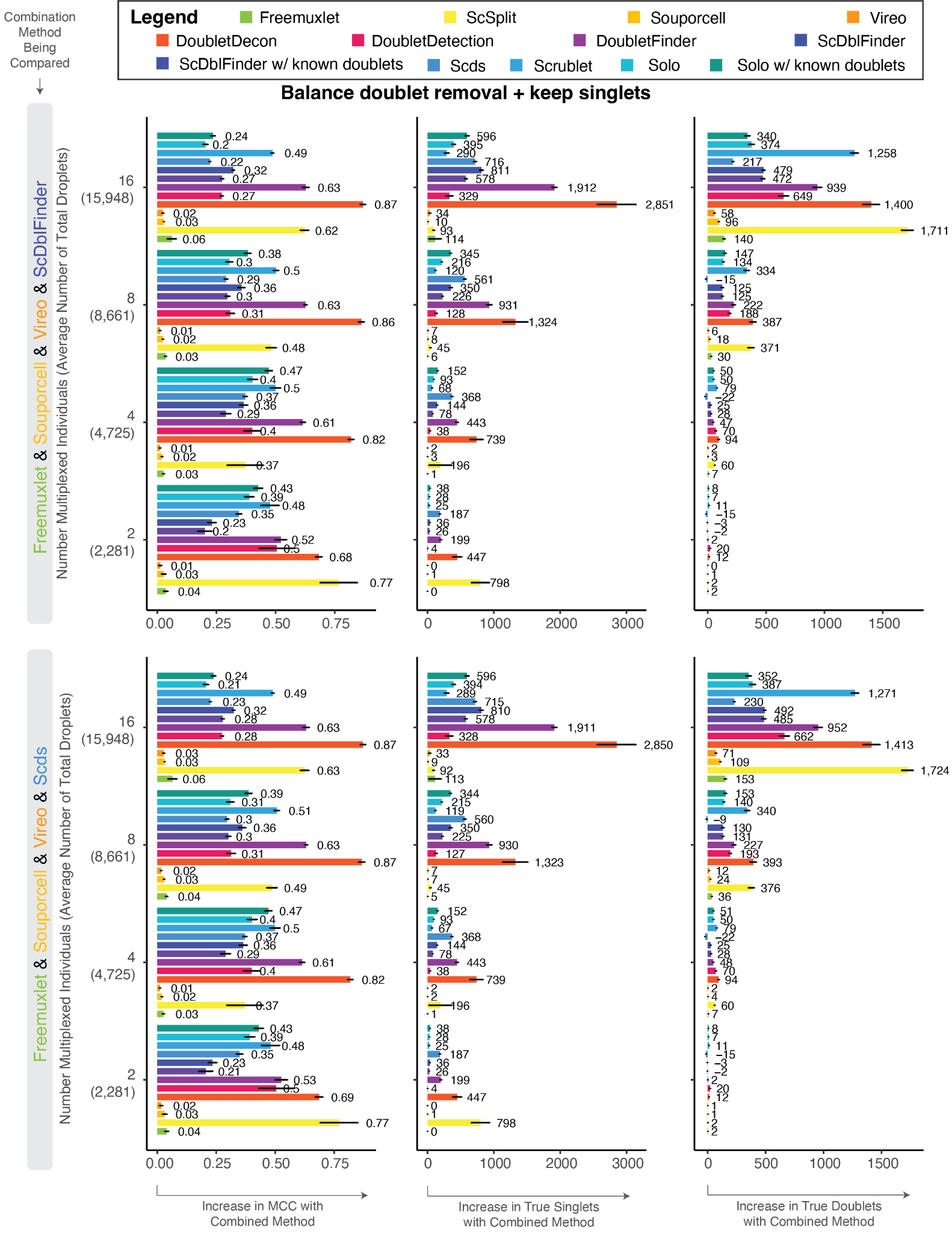

**Supplementary Figure 17: Change in MCC, True Singlets and True Doublets with Intersectional Methods Compared to Individual Methods for Multiplexed Pools with 16 or Fewer Individuals without reference SNP Genotypes. The average** increase in the MCC (left) and the resulting changes in true singlets (middle) and true doublets (right) for the recommended intersectional methods compared to the individual techniques for multiplexed pools with 16 or fewer individuals when reference SNP genotypes are unavailable when aiming to balance accurate doublet removal and maintaining true singlets. Numbers are represented as averages of method performance across ten pools for each multiplexed pool number. Error bars represent the standard error around the average across the ten pools for each pool size.

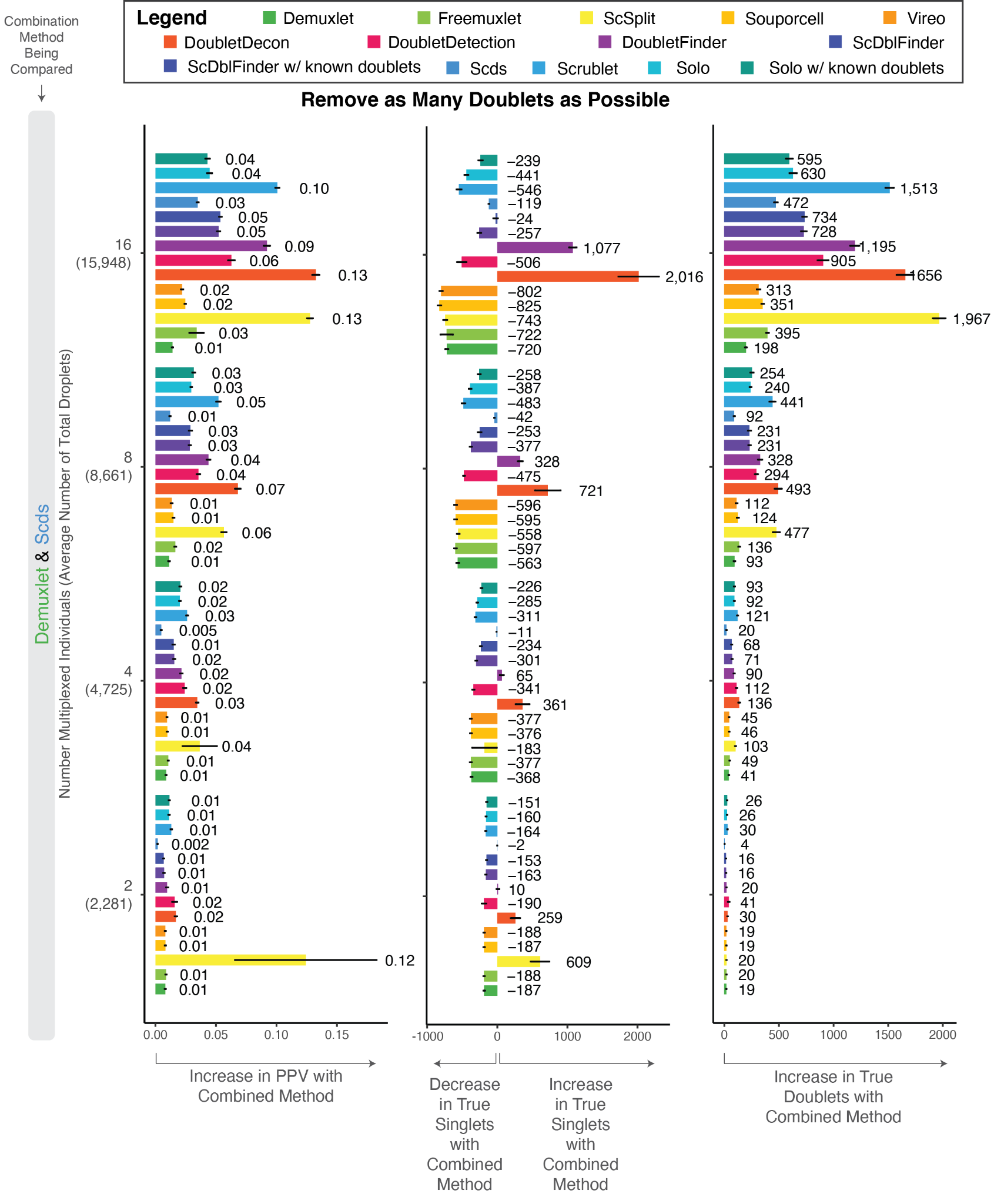

**Supplementary Figure 18: Change in PPV, True Singlets and True Doublets with Intersectional Methods Compared to Individual Methods for Multiplexed Pools with 16 or Fewer Individuals. The average** increase in the PPV (left) and the resulting changes in true singlets (middle) and true doublets (right) for the recommended intersectional methods compared to the individual techniques for multiplexed pools with 16 or fewer individuals when aiming to remove as many false singlets as possible. Numbers are represented as averages of method performance across ten pools for each multiplexed pool number. Error bars represent the standard error around the average across the ten pools for each pool size.

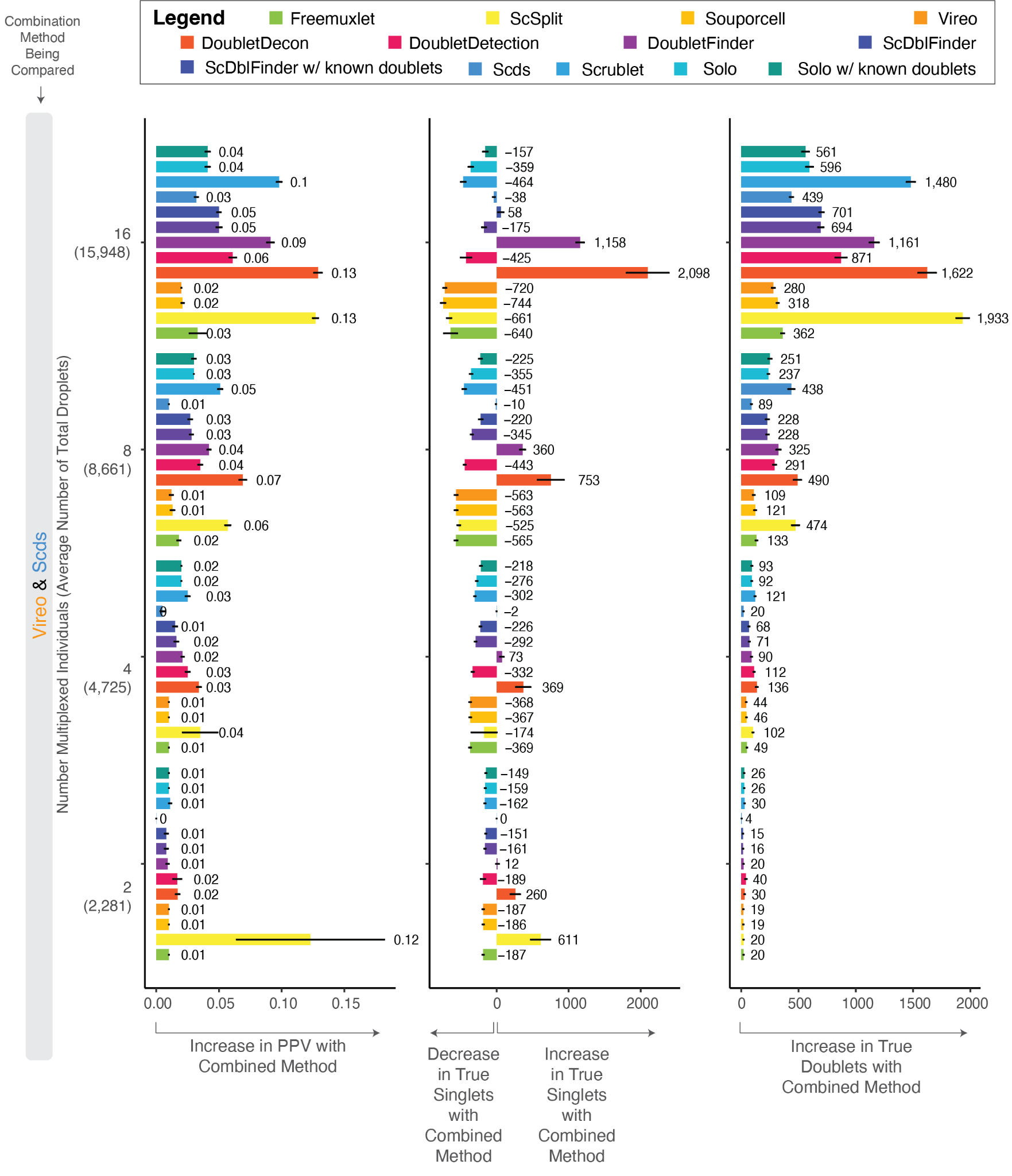

**Supplementary Figure 19: Change in PPV, True Singlets and True Doublets with Intersectional Methods Compared to Individual Methods for Multiplexed Pools with 16 or Fewer Individuals without Reference SNP Genotypes. The average** increase in the PPV (left) and the resulting changes in true singlets (middle) and true doublets (right) for the recommended intersectional methods compared to the individual techniques for multiplexed pools with 16 or fewer individuals when reference SNP genotypes are unavailable to remove as many false singlets as possible. Numbers are represented as averages of method performance across ten pools for each multiplexed pool number. Error bars represent the standard error around the average across the ten pools for each pool size.

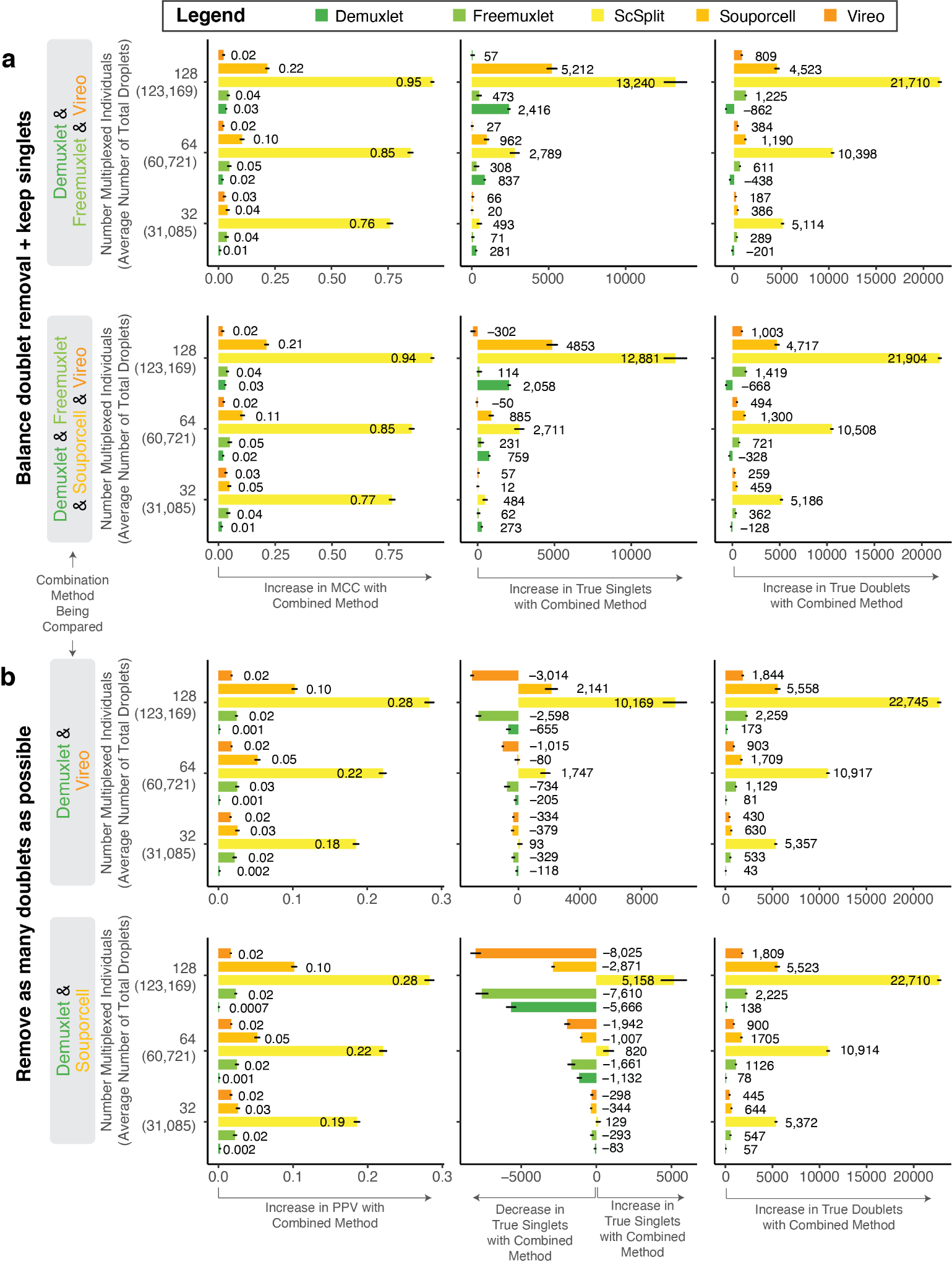

**Supplementary Figure 20: Change in MCC, PPV, True Singlets and True Doublets with Intersectional Methods Compared to Individual Methods for Large Multiplexed Pools. a)** Average increase in the MCC (left) and the resulting changes in true singlets (middle) and true doublets (right) for recommended intersectional methods compared to individual methods for large multiplexed pools when aiming to use a balanced method to remove true doublets but not at the expense of removing too many true singlets. **b)** Increase in the PPV (left) and the resulting changes in true singlets (middle) and true doublets (right) for recommended intersectional methods compared to individual methods for large multiplexed pools when aiming to remove as many false singlets as possible. Numbers are represented as averages of method performance across ten pools for each multiplexed pool number. Error bars represent the standard error around the average across the ten pools for each pool size.

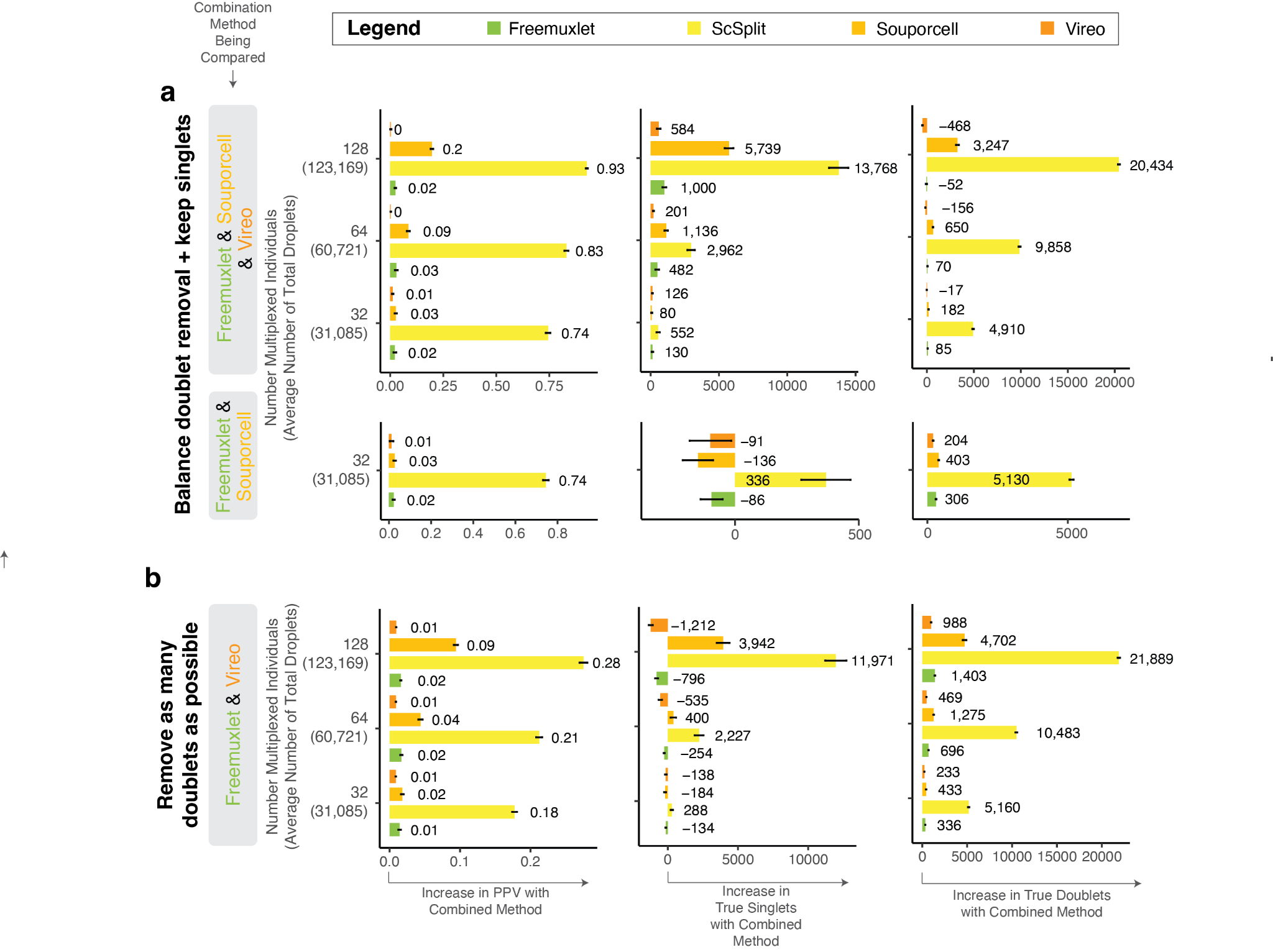

**Supplementary Figure 21: Change in MCC, PPV, True Singlets and True Doublets with Intersectional Methods Compared to Individual Methods for Large Multiplexed Pools without Reference SNP Genotypes. a)** Average increase in the MCC (left) and the resulting changes in true singlets (middle) and true doublets (right) for recommended intersectional methods compared to individual methods for large multiplexed pools without reference SNP genotypes when aiming to use a balanced method to remove true doublets but not at the expense of removing too many true singlets. **b)** Increase in the PPV (left) and the resulting changes in true singlets (middle) and true doublets (right) for recommended intersectional methods compared to individual methods for large multiplexed pools without reference SNP genotypes when aiming to remove as many false singlets as possible. Numbers are represented as averages of method performance across ten pools for each multiplexed pool number. Error bars represent the standard error around the average across the ten pools for each pool size.

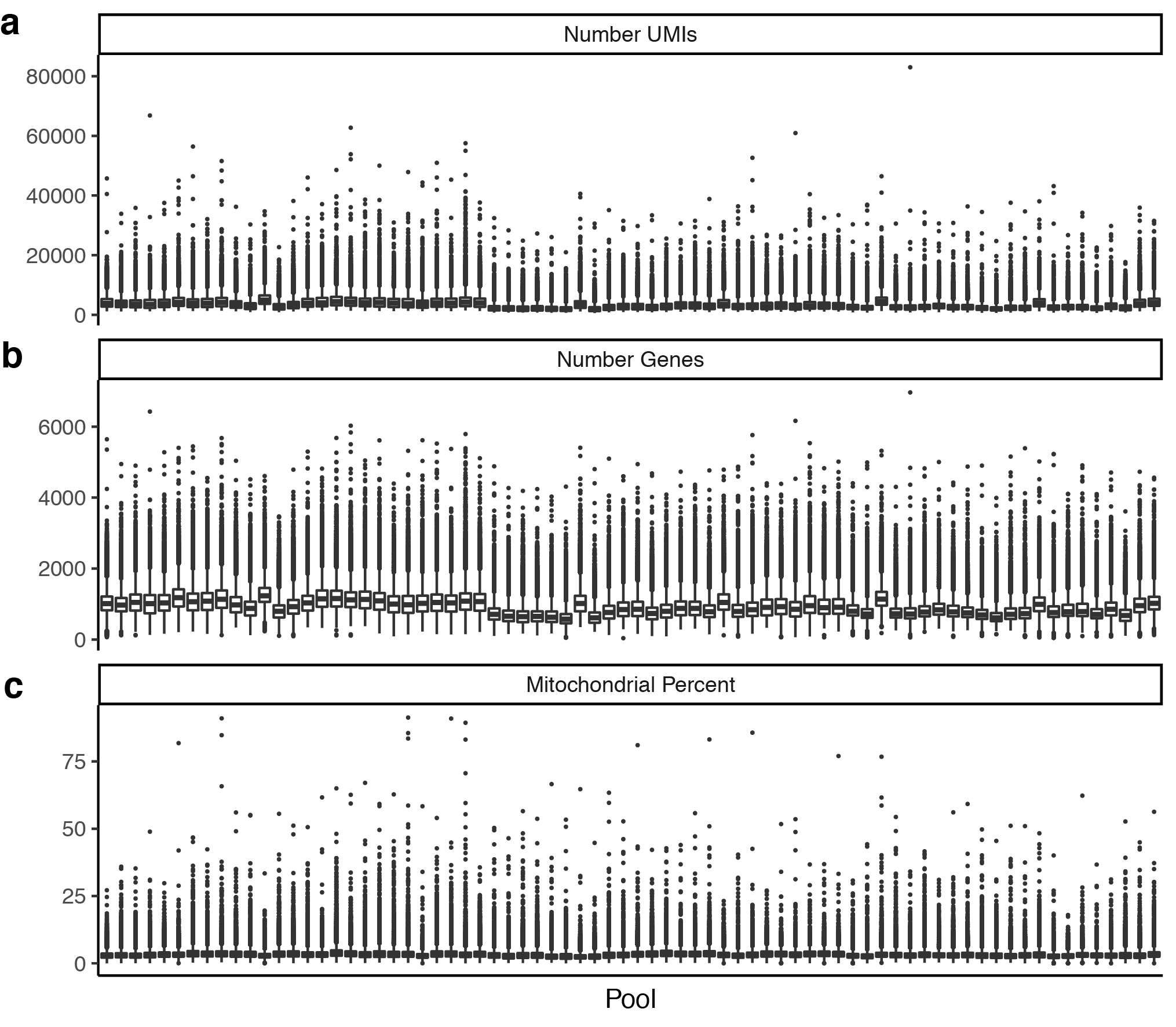

**Supplementary Figure 22: Quality Control Metrics of the PBMC pools.** The number of UMIs (**a**), number of genes (**b**) and mitochondrial per cent (**c**) per pool for the peripheral blood mononuclear cell (PBMC) dataset.

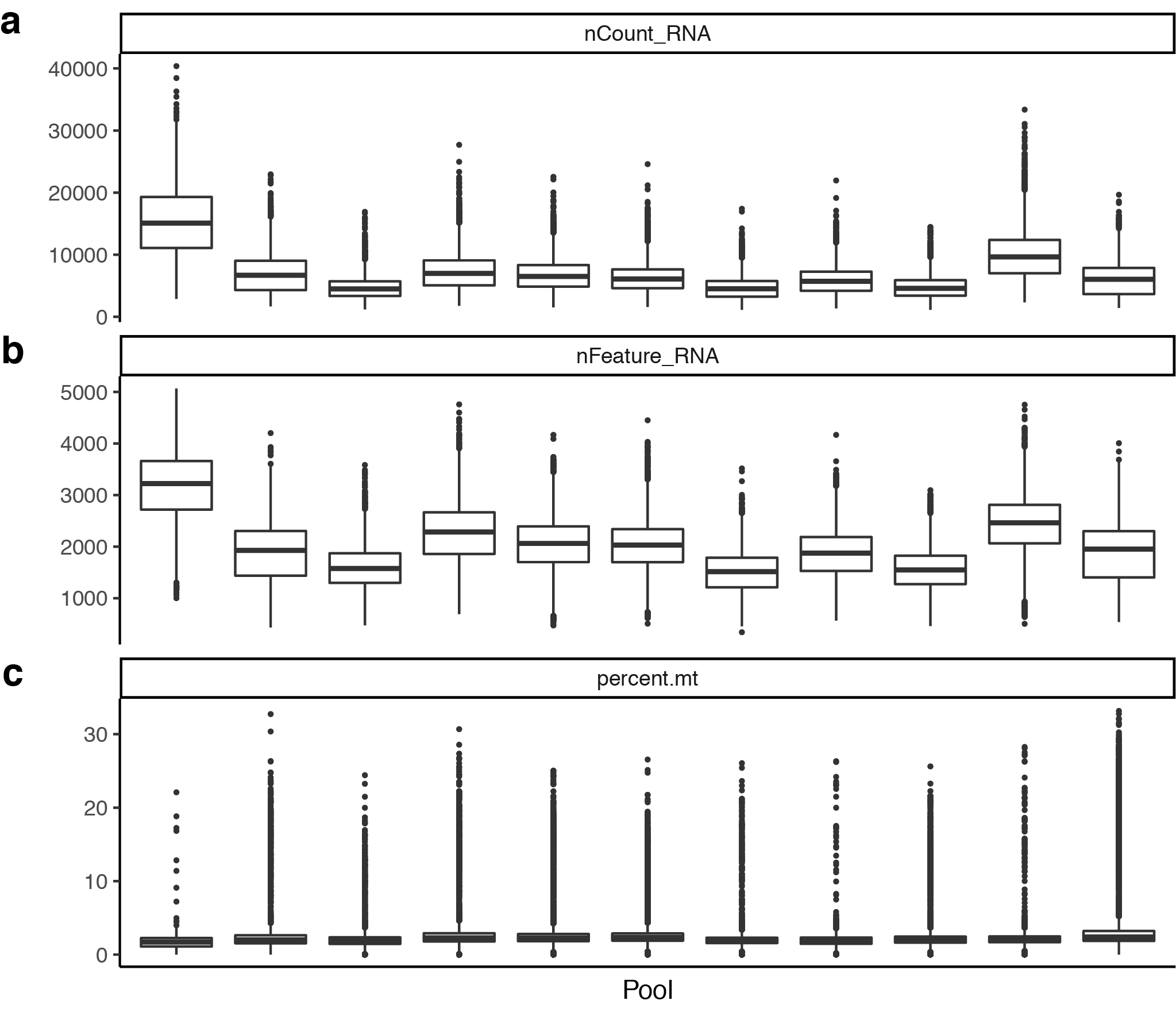

**Supplementary Figure 23: Quality Control Metrics of the fibroblast pools.** The number of UMIs (**a**), number of genes (**b**) and mitochondrial per cent (**c**) per pool for the fibroblast dataset.
